# Coordination of KIF3A and KIF13A regulates leading edge localization of MT1-MMP to promote cancer cell invasion

**DOI:** 10.1101/2021.05.24.445438

**Authors:** Valentina Gifford, Anna Woskowicz, Noriko Ito, Stefan Balint, Michael L. Dustin, Yoshifumi Itoh

## Abstract

MT1-MMP plays a crucial role in promoting the cellular invasion of cancer cells by degrading the extracellular matrix to create a path for migration. During this process, its localization at the leading edge of migrating cells is critical, and it is achieved by targeted transport of MT1-MMP-containing vesicles along microtubules by kinesin superfamily proteins (KIFs). Here we identified three KIFs involved in MT1-MMP vesicle transport: KIF3A, KIF13A, and KIF9. Knockdown of KIF3A and KIF13A effectively inhibited MT1-MMP-dependent collagen degradation and invasion, while knockdown of KIF9 increased collagen degradation and invasion. Our data suggest that KIF9 competes with KIF3A/KIF13A to bring MT1-MMP vesicles to different locations in the plasma membrane. Live-cell imaging analyses have indicated that KIF3A and KIF13A coordinate to transport the same MT1-MMP-containing vesicles. Taken together, we have identified a unique interplay between three KIFs to regulate leading edge localization of MT1-MMP and MT1-MMP-dependent cancer cell invasion.

## INTRODUCTION

Invasion and metastasis are the life-threatening features of invasive cancer. Epithelial cancer cells achieve this by losing their cell-cell adhesion property, increasing their motility, and gaining extracellular matrix (ECM) degrading activity for invasion. It has been shown that one of the crucial ECM-degrading enzymes allowing cancer cell invasion and metastasis is membrane-type I matrix metalloproteinase, MT1-MMP(Itoh, 2015; Sabeh et al., 2004; Sabeh et al., 2009; Sato et al., 1994; Seiki, 2003). MT1-MMP is a type-I transmembrane proteinase that belongs to the matrix metalloproteinase (MMP) family. MT1-MMP degrades various ECM components on the cell surface, including fibrillar collagen (Holmbeck et al., 2004; Itoh, 2015; Ohuchi et al., 1997). MT1-MMP also activates proMMP-2 and proMMP-13 on the cell surface, expanding proteolytic repertoire(Itoh, 2015). ProMMP-2 activation is considered essential for epithelial cancer cell invasion and growth since activated MMP-2, but not MT1-MMP, can degrade type IV collagen, a major component of the basement membrane(Itoh, 2015; Taniwaki et al., 2007). MT1-MMP also cleaves membrane proteins, including CD44(Kajita et al., 2001), ICAM-1(Sithu et al., 2007), LRP1(Rozanov et al., 2004), syndecan 1(Endo et al., 2003), ADAM9(Chan et al., 2012), Dll1(Jin et al., 2011), EphA2(Koshikawa et al., 2015; Sugiyama et al., 2013), modifying cell adhesion property and cellular signalling. Thus, MT1-MMP is considered to be a microenvironment and cell function modifier.

Localization of MT1-MMP at the leading edge of migrating cells is crucial to promote cellular invasion(Ferrari et al., 2019; Gifford and Itoh, 2019; Itoh, 2015). Leading-edge structures or their precursors include filopodia, lamellipodia, invadopodia, and podosomes, and MT1-MMP localizes to all of these structure(Ferrari et al., 2019; Gifford and Itoh, 2019; Itoh, 2015). MT1-MMP also localizes at focal adhesion (FA)(Wang and McNiven, 2012; Woskowicz et al., 2013). It is thought that localization of MT1-MMP is achieved by direct transport of MT1-MMP-containing vesicles to these membrane structures(Wiesner et al., 2010), but the mechanism of vesicle transport of MT1-MMP is still poorly understood. Vesicle transport is carried out by motor proteins, including kinesin superfamily proteins (KIFs), which transport vesicles and macromolecules along microtubules(Hirokawa et al., 2009; Hirokawa and Tanaka, 2015). There are 45 KIFs in humans that are classified into three groups, N-, M- and C-kinesins, according to the position of the microtubule-binding motor domain(Hirokawa and Tanaka, 2015). KIFs that transport vesicles toward the cell periphery or the (+) ends of microtubules belong to N-kinesins, which form the largest subgroup of 39 KIFs and can be further divided into 11 subgroups(Hirokawa et al., 2009; Hirokawa and Tanaka, 2015). N-kinesins have a motor domain at their N-terminus, followed by a neck region, a coiled-coil region, and a C-terminal tail region(Hirokawa et al., 2009; Hirokawa and Tanaka, 2015). It is thought that each KIF can selectively recognize certain cargos through the specific interaction with adaptor molecules, membrane proteins, or Rab GTPases through their C-terminal tail region. So far, KIF5B and KIF3A/B have been reported to transport MT1-MMP vesicles in macrophages (Wiesner et al., 2010). KIF5B and KIF3A have been reported to play a role in invadopodia localization of MT1-MMP in MDA-MB231 cells (Marchesin et al., 2015; Thapa and Anderson, 2017). However, the roles of KIFs in MT1-MMP vesicles trafficking in different cells are not understood.

This study aimed to identify KIFs responsible for MT1-MMP localization at the leading edge of invasive human fibrosarcoma, HT-1080. By screening with siRNAs, we identified three KIFs (KIF3A, KIF13A, and KIF9) directly involved in MT1-MMP vesicle trafficking. We report here the co-ordination of KIF3A and KIF13A to transport MT1-MMP vesicles and the potential competitive role of KIF9 against KIF3A and KIF13A.

## RESULTS

### Knockdown of KIF13A, KIF3A, KIF9 and KIF1C alters MT1-MMP-medited cell functions on the cell surface

To identify KIFs responsible for MT1-MMP vesicle transport, we initially narrowed down the candidate KIFs. There are 45 KIFs in humans, and we initially excluded C- and M-kinesins since they do not traffic to the (+) ends of microtubules. We further excluded KIFs reported to be exclusively expressed in neurons or involved in cell division as they are unlikely to control MT1-MMP vesicle trafficking (Supplementary Table S1). This exercise leaves 17 KIFs, including splicing variants, as candidates for screening. In order to screen KIFs necessary for MT1-MMP trafficking, we set up a gelatin and collagen film degradation assay with HT-1080 cells. As shown in Supplementary Figure S1a-d, gelatin and collagen film degradation were abolished by a broad-spectrum MMP inhibitor GM6001 (10 μM) and by a specific biologic MT1-MMP inhibitor DX-2400 (200nM)(Devy et al., 2009), but not by TIMP-1 (200 nM), which does not inhibit MT1-MMP but inhibits all soluble and GPI-anchored MMPs. The data suggested that both gelatin and collagen film degradation activities are solely dependent on endogenous MT1-MMP in HT1080 cells. We confirmed that HT-1080 cells expressed all 17 KIFs (Supplementary Fig S2a), each KIF gene was silenced by siRNA, and cells were subjected to gelatin film degradation assay (Figure S2b, c). Among the 17 KIFs, knockdown of KIF1C, KIF3A, and KIF13A notably decreased gelatin film degradation, while knockdown of KIF9 rather increased gelatin film degradation (Figure S2b). Therefore, we investigated KIF1C, 3A, 13A and 9 further.

Knockdown of KIF13A, KIF3A, and KIF1C notably decreased MT1-MMP-mediated gelatin film degradation, whereas the knockdown of KIF9 enhanced it in a statistically significant manner compared to non-targeting siRNA (si-NT) transfected cells (P < 0.0001) (Figure 1a). Western Blotting for KIF3A and KIF1C and by RT-PCR for KIF13A and KIF9 (Figure 1d, top and middle panels) confirmed the efficiency of silencing. Interestingly, siRNA for KIF9 by smart pool siRNA (si-KIF9) was selective for KIF9-v1 (knocked down by more than 90%), as KIF9-v2 and −3 mRNA level were knocked down only by 39%. We also confirmed that silencing any of these *kinesin* genes did not alter MT1-MMP mRNA levels (Figure 1d, bottom panel). Since it has been reported that KIF5B mediates MT1-MMP intracellular trafficking in primary macrophages and breast cancer cell line, MDA-MB231(Marchesin et al., 2015; Thapa and Anderson, 2017; Wiesner et al., 2010), we also re-examined KIF5B knockdown in HT-1080 cells. We confirmed that KIF5B knockdown did not affect gelatin film degradation (Figure 1e). Next, we examined the effect of KIF knockdown on the collagenolytic activity of MT1-MMP in HT-1080 cells as the type I collagen represents its physiological substrate. As shown in Figure 1b, KIF13A, KIF3A, and KIF1C knockdowns decreased collagen degradation by HT1080 cells, while the knockdown of KIF9 enhanced it in a statistically significant manner (Figure 1b, right panel).

**Figure 1.**
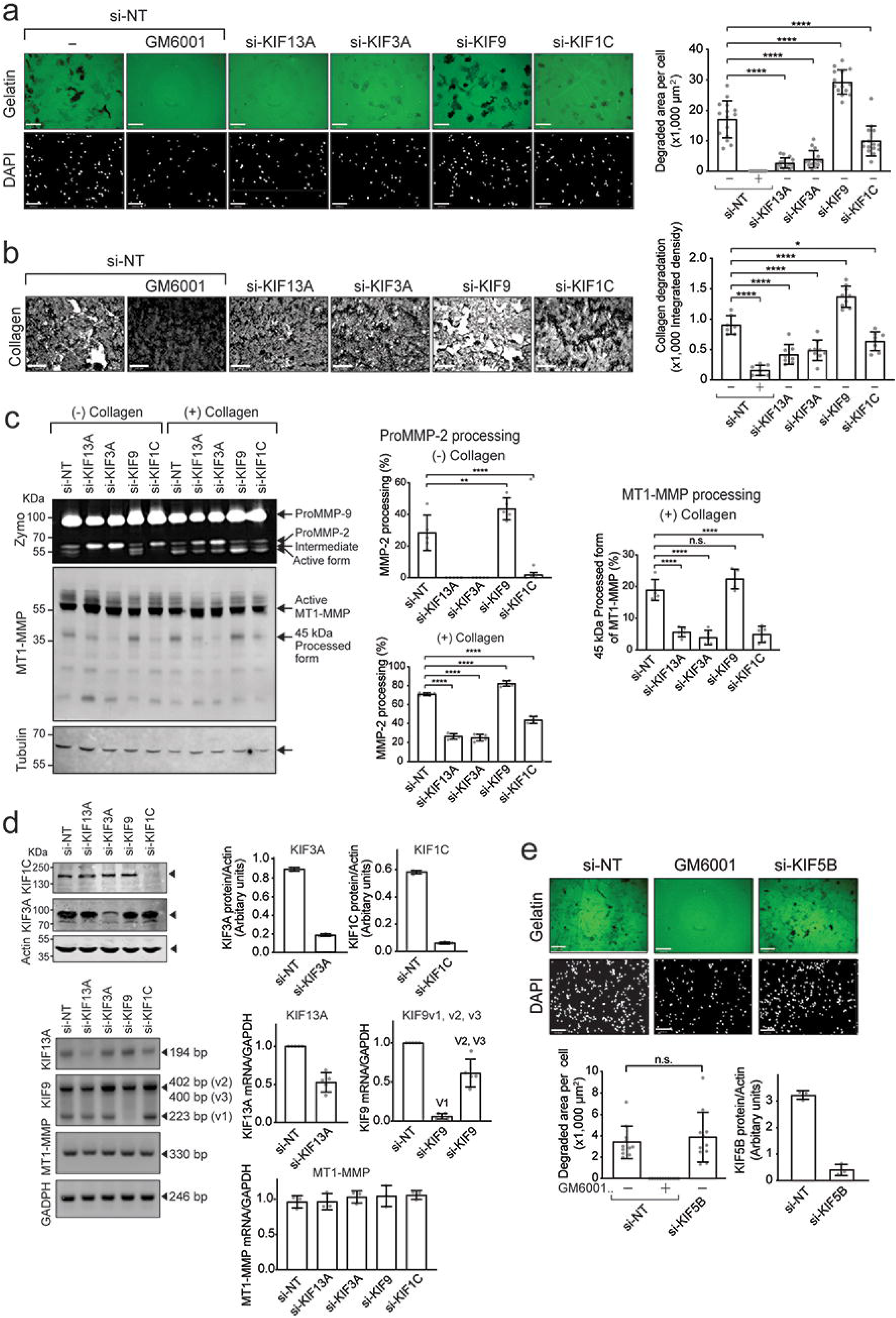
Knockdown of KIF13A, KIF3A, KIF9 and KIF1C alters MT1-MMP-mediated cell functions on the cell surface. **a.** HT-1080 cells were transfected with the indicated siRNAs and subjected to gelatin film degradation assay in the presence or absence of GM6001 (10 μM). Scale bars are 130 μm. Quantification of the degradation area (μm^2^) per cell in HT-1080 cells transfected with the indicated siRNAs was show as a graph (right panel). Data are presented as the mean of fifteen independent microscopic fields of view and are representative of five independent experiments. One-way ordinary Anova with Tuckey’s multiple comparisons test. Data are shown as mean ± SD. \*\*\*\**P* < 0.0001. **b.** HT-1080 cells were transfected with the indicated siRNAs and subjected to collagen film degradation assay in the presence or absence of GM6001 (10 μM). Scale bars are 130 μm. Quantification of the integrated density of the collagen layer in HT-1080 cells transfected with the indicated siRNAs was shown as a graph (right panel). Data are presented as the mean of eight independent microscopic fields of view and are representative of five independent experiments. One-way ordinary Anova with Tuckey’s multiple comparisons test. Data are shown as mean ± SD. \*\*\*\**P* < 0.0001; \**P* < 0.05. **c.** HT-1080 cells transfected with the indicated siRNAs were cultivated in the presence or absence of collagen I (100 μg/ml) for 24 h. Culture media were analysed by zymography (Zymo) for proMMP-2 activation. Cell lysates were subjected to Western blotting using antibodies for MT1-MMP Hpx domain and tubulin. Arrows point bands of interest. ProMMP-2, 68 kDa proMMP-2; Intermediate, 65 kDa intermediate form; Active form, 62 kDa active MMP-2. Quantification of the percentage of MMP-2 processed forms over the total was shown as a graph (middle panel). Top panel shows processed MMP-2 without collagen stimulation, and the bottom panel shows the MMP-2 activation upon collagen stimulation. Quantification of the 45 kDa MT1-MMP processed forms (%) upon collagen stimulation was shown as a graph (right panel). Data are calculated from five independent experiments. One-way ordinary Anova with Tuckey’s multiple comparisons test. Data are presented as mean of five blots representative of five independent experiments. One-way ordinary Anova with Tuckey’s multiple comparisons test. Data are shown as mean ± SD. \*\*\*\**P* < 0.0001; \*\**P* < 0.01. n.s., non-significant. **d.** Efficiency of KIF knockdown was assessed by Western blotting for KIF1C and KIF3A and by RT-PCR for KIF13A, KIF9v1-v3. Effect of KIF knockdown on MT1-MMP mRNA level was also examined. Quantification data were shown as a graph. Western blot band intensities were normalized with Actin band intensities and mRNA normalized with the one for GAPDH. **e**. HT1080 cells transfected with siRNA for KIF5B were subjected to gelatin film degradation assay. Top panel shows representative images from each treatment as indicated. Bottom left panel shows a quantitative data as degradation area per cells. Data is shown as mean ± SD (n=12). n.s., non-significant. Bottom right panel shows a quantitative data of KIF5B measured by Western blot. Data are presented as mean of three blots representative of three independent experiments.

Another cell surface activity of MT1-MMP is proMMP-2 activation. ProMMP-2 activation can be induced by adding collagen I (100 μg/ml) in the culture medium(Majkowska et al., 2017). We confirmed that the collagen-induced proMMP-2 activation is due to MT1-MMP in HT-1080 cells (Supplementary Figure S1e, f). As indicated in Figure 1c, the conditioned medium from si-NT-transfected cells, without collagen, showed proMMP-2 (ProMMP-2, 68 kDa) and its intermediate form (Intermediate, 65 KDa) (Figure 1c, left panel). Silencing *Kif13a, Kif3a*, and *Kif1c* genes prevented the generation of the intermediate form, whereas silencing the *Kif9* gene accelerated proMMP-2 processing to its intermediate and active forms (Active form, 62 kDa, Figure 1c, left panel). Upon collagen stimulation, around 71% of proMMP-2 was converted to its active form in si-NT-transfected cells (Figure 1c, center lower panel). The knockdown of KIF13A and KIF3A significantly reduced the activation down to 26% and 25%, respectively. Knockdown of KIF1C seemed to moderately decrease it to 44% (Figure 1c, center lower panel). Conversely, silencing the *Kif9* gene enhanced the activation to around 80% in a statistically significant manner (Figure 1c, center lower panel). Interestingly, we noted that the generation of the 45 KDa processed form of MT1-MMP, which has been shown to coincide with functional activation of MT1-MMP, was also affected by KIF knockdown (Figure 1c, left panel)(Stanton et al., 1998). The knockdown of KIF13A, KIF3A, and KIF1C significantly reduced the generation of the 45 kDa form (Figure 1c, left and right panels), while the knockdown of KIF9 have a tendency to increase it. However, the effect was statistically not significant (Figure 1c, left and right panels).

### Knockdown of KIF13A, KIF3A, KIF9 and KIF1C alters MT1-MMP localization at the cell-matrix interface

The cell surface biotinylation experiment indicated that KIF knockdown did not affect the overall cell surface level of MT1-MMP (Figure 2a). This result was also confirmed by immunofluorescence staining of the cell surface MT1-MMP (Figure 2b). Therefore, we hypothesized that the KIF knockdown shifts the localization of MT1-MMP to or from the cell-matrix interface. We employed total internal reflection fluorescent (TIRF) microscopy to examine the levels of MT1-MMP at the ventral side of the membrane. As shown in Figure 2c, silencing *Kif13a* and *Kif3a* genes significantly reduced MT1-MMP localization at the cell-matrix interface, whereas the knockdown of KIF9 increased it (Figure 2c). Knockdown of KIF1C did not change MT1-MMP localization significantly (Figure 2c). The knockdown of these KIFs did not influence cellular attachment to the gelatin, as it did not impact the level of β1 integrin (Figure 2d) and close cell-matrix contacts detected by interference reflection microscopy (IRM) (Figure 2d, top panel). Taken together, these data indicate that knocking down KIF3a, KIF13a and KIF9 altered MT1-MMP-dependent activity by changing the level of MT1-MMP localization at the cell-matrix interface.

**Figure 2.**
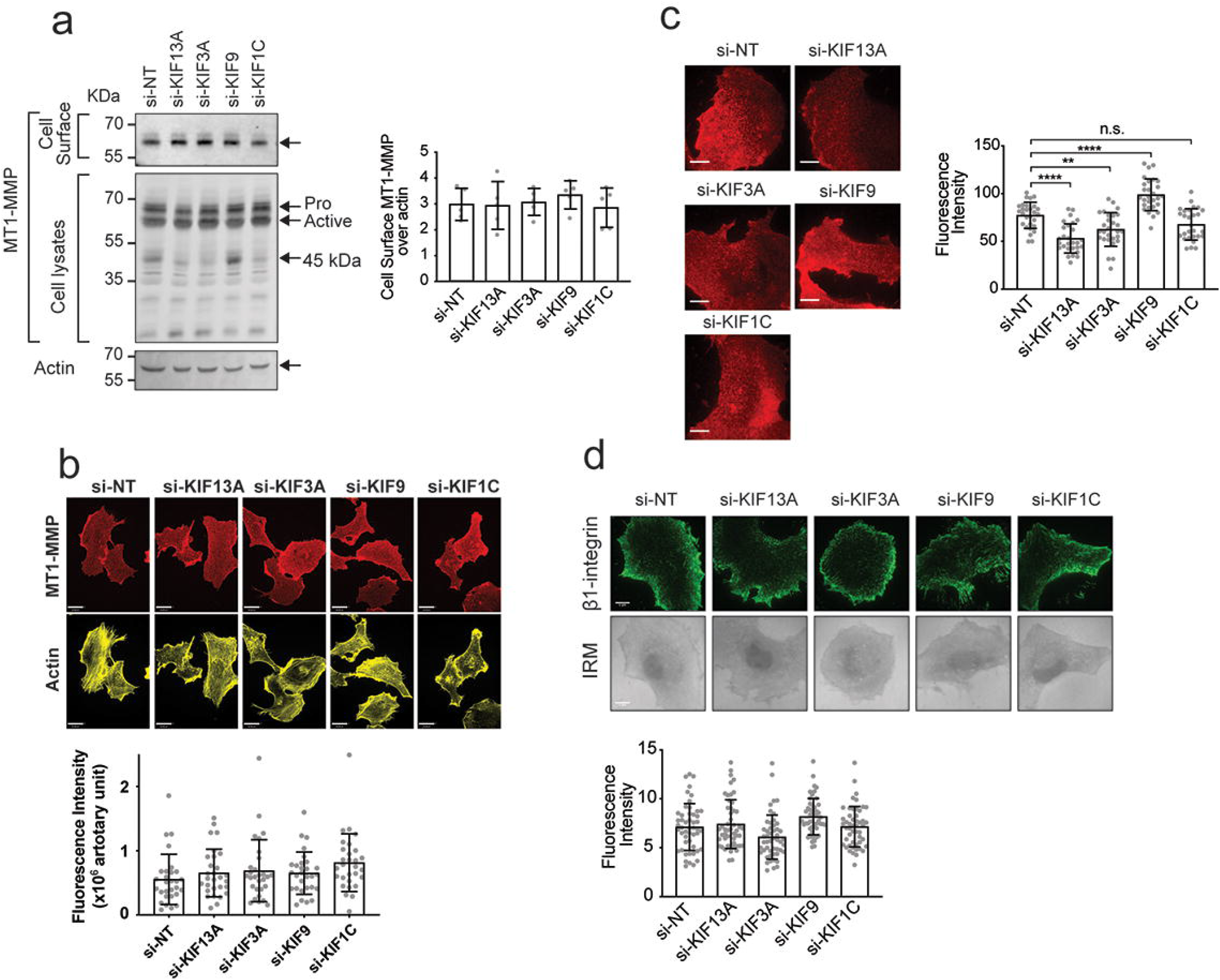
Knockdown of KIF13A, KIF3A, KIF9 and KIF1C modifies MT1-MMP localisation at the substrate-attachment side. **a.** HT-1080 cells were transfected with the indicated siRNAs subjected to cell surface biotinylation. Proteins eluted from the streptavidin beads were analysed by Western blot using anti-Loop MT1-MMP antibody. Whole cell lysates were analysed by Western blot using 1D8 anti-MT1-MMP and anti-actin antibodies. Arrows point bands of interest. Quantification of the cell surface MT1-MMP in the left panel is shown as a graph (right panel). MT1-MMP band intensities were normalized by actin band. Data are presented as mean ± SD. (n=5) **b.** HT1080 cells transfected with siRNAs targeting KIFs were subjected to indirect immunofluorescent staining with anti-MT1-MMP antibody without permeabilization and imaged with confocal microscopy. Representative extended focus images of three independent experiments were shown for MT1-MMP (red) and actin (yellow) (top panel). Quantitative data were shown as a graph (bottom panel). Fluorescent intensity of extended focus image per cell was measured and the data are presented as mean ± SD. **c.** HT-1080 cells transfected with siRNAs targeting the KIFs were stained with anti-MT1-MMP antibody without permeabilization (red) and imaged by TIRF microscopy. Representative images of three independent experiments are shown (left panel). Scale bars are 10 μm. Quantitative data of the fluorescent intensities at the substrate-attachment side were shown as a graph (right panel). Data are presented as mean ± SD. (n=30). \*\*\*\**P* < 0.0001; \*\**P* < 0.01; n.s., non-significant. **d.** HT-1080 cells were transfected with siRNAs targeting the selected KIFs. After 72 hours, cells were stained with anti-β1 integrin (green) antibody without permeabilisation and imaged by TIRF microscopy. Representative images of three independent experiments are shown (top panel). Scale bars are 10 μm. Quantification of β1 integrin expression at the substrate attachment side in HT-1080 cells were shown as a graph (bottom panel). Data are presented as mean ± SD. (n=30).

### The knockdown of KIF13A, KIF3A and KIF9 alters MT1-MMP-mediated cell invasion through a microporous membrane and MT1-MMP-dependent cell migration in 3D collagen

We next examined the effect of KIF knockdown on MT1-MMP-dependent cell invasion. We employed two different invasion assays: collagen invasion assay using trans-well chambers coated with collagen gel (Figure 3a) and microcarrier bead invasion assay that assesses migration distance of cells from the beads’ surface within 3D collagen gel (Figure 3b). The cellular invasion in these assays is entirely dependent on endogenous MT1-MMP in HT-1080 cells (Figure S1f, g). As shown in Figure 3a (bottom right panel), the knockdown of KIF13A and KIF3A significantly decreased the invasion, whereas KIF9 knockdown significantly enhanced it in the trans-well invasion assay. KIF1C knockdown did not affect the cell invasion (Figure 3a, bottom right panel). In microcarrier bead invasion assay (Figure 3b), the knockdowns of KIF13A, KIF3A, and KIF1C significantly reduced cell migration. Interestingly, si-KIF9-transfected cells also migrated significantly less (Figure 3b), which is contrary to the trans-well invasion assay (Figure 3a). The effect of knockdown of KIF13A, KIF3A, and KIF9 on cellular invasion was not due to alteration on the fundamental cell migration machinery. These cells migrated similarly on the plastic surface as measured by a wound-healing assay (Figure 3c). On the other hand, KIF1C knockdown slightly decreased it (P = 0.019), indicating that KIF1C may be involved in the general cell migration (Figure 3c).

**Figure 3.**
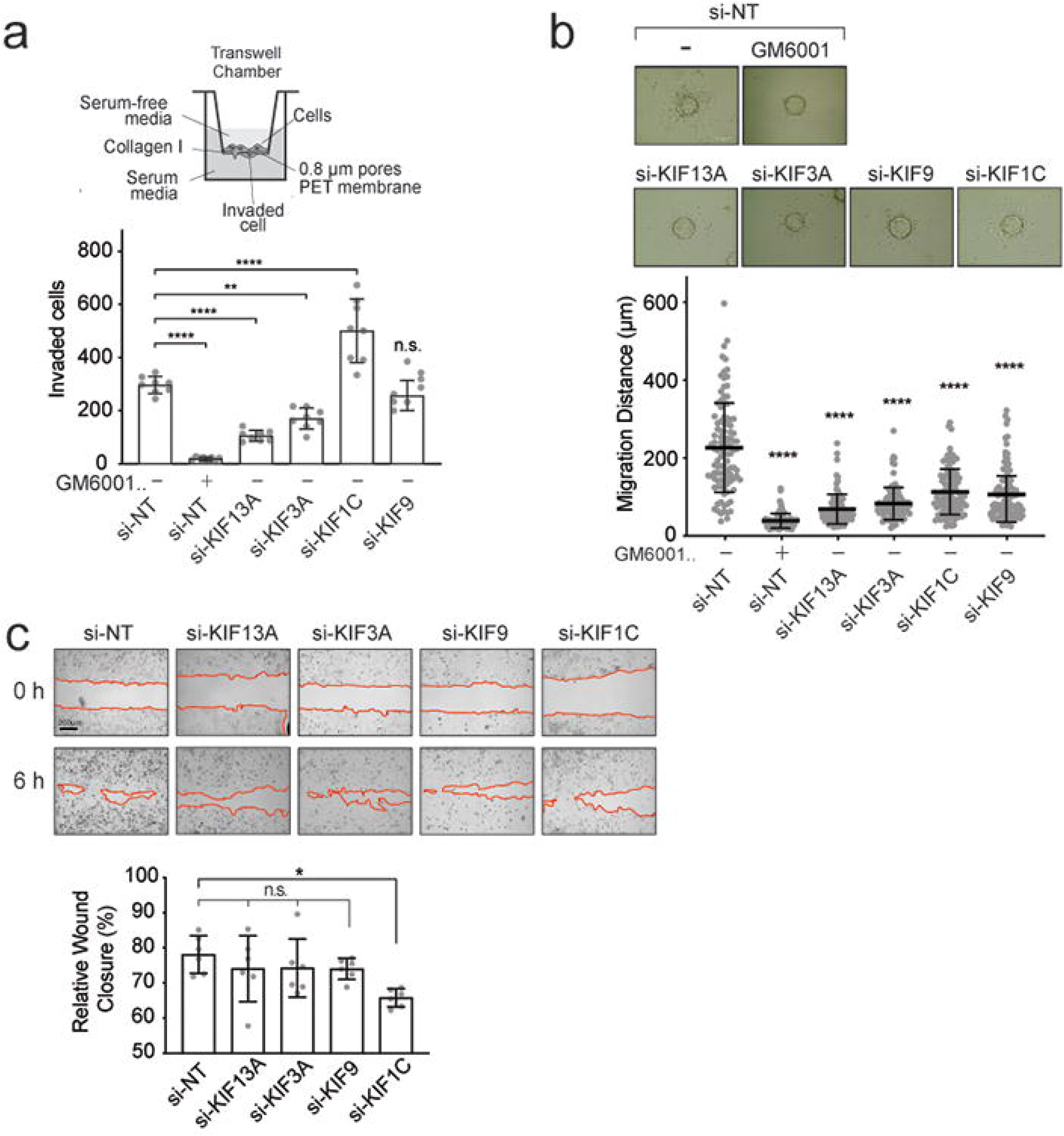
The knockdown of KIF13A, KIF3A, KIF9 and KIF1C diminishes MT1-MMP-dependent cell migration in 3D collagen. **a.** HT1080 cells transfected with siRNA indicated were subjected to the same invasion assay and data shown as a graph (bottom right panel). Data are presented as mean of the number of cells migrated through six trans-wells for each treatment and are representative of three independent experiments. **** *P* < 0.0001, ** *P* < 0.001, n.s., non-significant. **b.** HT1080 cells were transfected with siRNAs targeting the selected KIFs and subjected to bead invasion assay in the presence or absence of GM6001 (10 μM). A representative image of cells migrating from the surface of the bead into collagen type-I matrix (2mg/ml) is shown (top panel). Scale bar,130 μm. Data are presented as mean of the distance migrated by one hundred cells for each treatment. Error bars SD, **** *P* < 0.0001, n.s., non-significant. **c.** HT1080 cells were transfected with siRNAs targeting the selected KIFs and subjected to wound healing assay on plastic. Images of 0h and 6h time points are shown. Images are representative of three independent experiments. Red lines highlight the migration front. Scale bar, 200 μm. Quantification of wound healing assay data were shown as a graph (bottom panel). Data are presented as mean of the percentage of wound closure relative to the initial gap area. * *P* < 0.019.

### KIF13A and KIF3A co-localize with MTI-MMP-containing vesicles

Next, the cellular distribution of MT1-RFP and GFP-tagged KIF13A, KIF3A, KIF9-v1, KIF9-v2, and KIF1C were analyzed. As shown in Figure 4a, KIF13A-GFP co-localized with MT1-RFP, and the two proteins exhibited a similar distribution pattern within the perinuclear (white box) and the cell periphery areas (orange box). KIF3A-GFP also co-localized with MT1-RFP-positive vesicles within the perinuclear region (Figure 4b, white box), but the two proteins did not co-localize at the cell periphery (Figure 4b, orange box). KIF9-v1-GFP co-localized with MT1-RFP within the perinuclear region (Figure 4c, White box), but not cell periphery (Figure 4c, orange box). KIF9-v2-GFP- and KIF1C-GFP-transfected cells did not show a clear vesicular-like distribution of MT1-RFP clearly, and neither KIF9-v2-GFP nor KIF1C-GFP showed co-localization with MT1-RFP neither in perinuclear nor cell periphery region (Figure 4d and e). It was interesting to note that KIF1C-GFP strongly accumulated at the tips of the trailing edges (Figure 4e, orange box). Pearson’s correlation coefficients (PCCs) confirmed that KIF13A-GFP and KIF3A-GFP had the highest degree of co-localization (Figure 4f). On the other hand, KIF9-v2-GFP and KIF1C-GFP exhibited the least co-localization with MT1-RFP. These cells were also embedded in a 3D collagen matrix and imaged by confocal microscopy (Figure 4g-i). The data confirmed that KIF13A-GFP exhibited the highest co-localization with MT1-RFP (Figure 4g, i), whereas KIF3A-GFP co-localization was found only within the perinuclear region (Figure 4h, i). We confirmed no co-localization of KIF9-v2-GFP and KIF1C-GFP with MT1-RFP under these conditions (Figure 4j-i).

**Figure 4.**
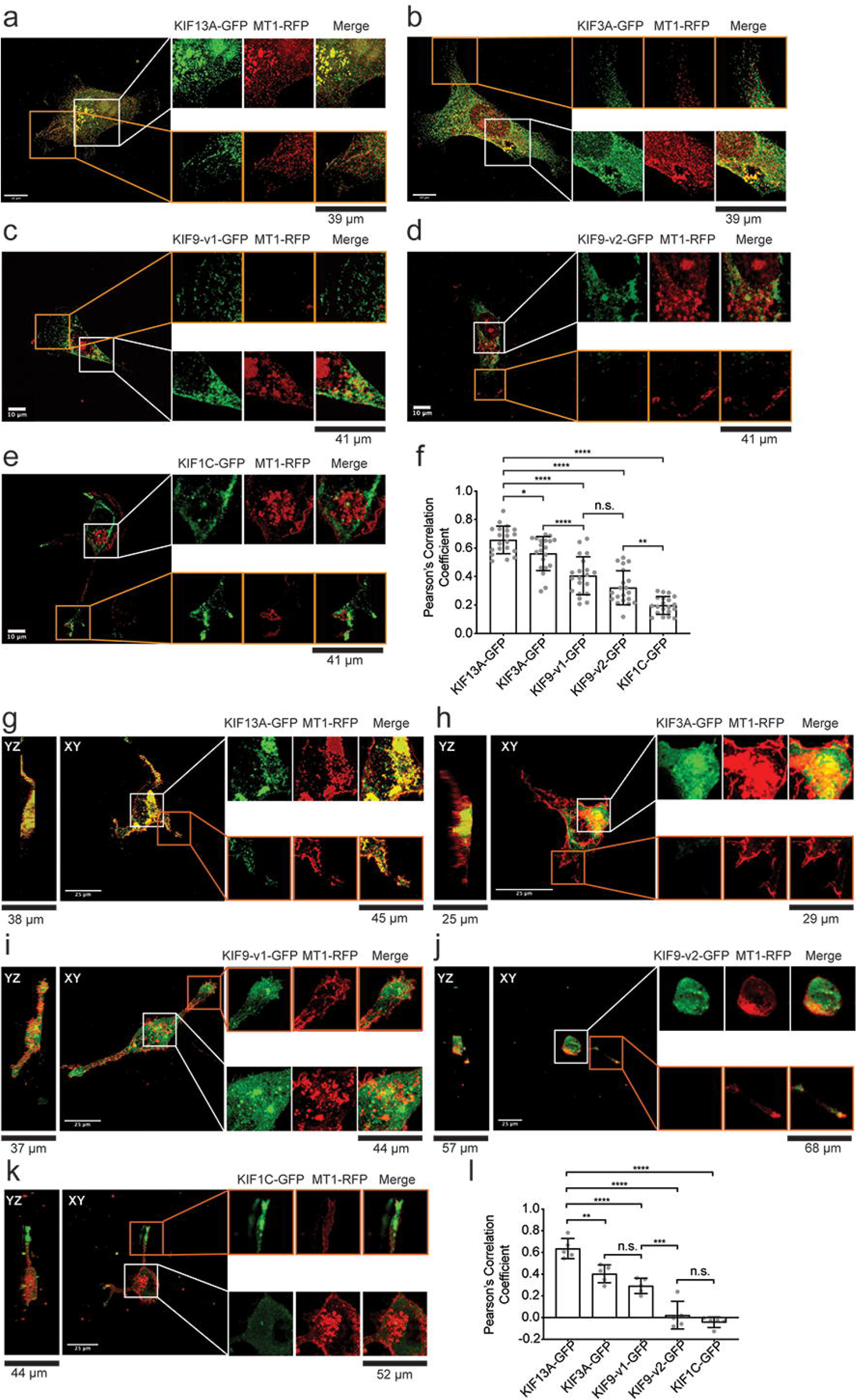
MT1-RFP and KIF-GFP co-localisation in HT-1080 cells cultured on gelatin-coated cover glass and within 3D collagen matrix. **a, b, c, d, e.** HT-1080 cells transfected with GFP-tagged KIFs (indicated, green) and MT1-RFP (red) were cultured on gelatin-coated cover glass. Extended focus representative images are shown. White boxes enclose enlarged areas for perinuclear region and orange boxes for periphery region. Scale bars indicate 10 μm. **f.** Quantification of co-localisation between GFP-tagged KIFs and MT1-RFP. Data are presented as the mean of 20 PCC calculated for 20 different cells per treatment and are representative of three independent experiments. One-way ordinary Anova with Tuckey’s multiple comparisons test. Data are shown as mean ± SD. \*\*\*\**P* < 0.0001: \**P* < 0.05; \*\**P* < 0.01; n.s., non-significant. **g, h, i, j, k.** HT-1080 cells transfected with GFP-tagged KIFs (green) and MT1-RFP (red) were cultured within 3D collagen gel and imaged by confocal microscopy. Extended focus representative images and the corresponding orthogonal views are shown. White boxes enclose enlarged areas. Scale bars are 25 μm. **l.** Quantification of co-localisation between GFP-tagged KIFs and MT1-RFP. Data are presented as the mean of five PCCs calculated for five different cells per treatment and are representative of three independent experiments. One-way ordinary Anova with Tuckey’s multiple comparisons test. Data are presented as mean ± SD; \*\**P* < 0.01; \*\*\**P* < 0.001; \*\*\*\**P* < 0.0001; n.s., non-significant.

### KIF13A, KIF3A and KIF9-v1 transport MT1-MMP-containing vesicles

These cells were next subjected to time-lapse imaging using confocal microscopy. The expression of KIF13A-GFP often made a distribution of MT1-RFP signals into a tubular-like shape, extending from the perinuclear regions towards the tips of the ruffling membrane (Figure 5a). Numerous KIF13A-GFP signals were observed moving along these MT1-RFP tubular-like structures, accumulating at the cell periphery, suggesting that the tubular-like structures are microtubules, and MT1-RFP seems to be distributed over the microtubules (supplemental *movie S1*). Vesicles double positive for KIF13A-GFP and MT1-RFP could also be detected within the perinuclear area. Despite MT1-RFP strongly accumulated within this region, it was possible to observe KIF13A-GFP transporting MT1-RFP-containing vesicles (Figure 5b, *movie S2*). No significant differences were observed in velocity or size of vesicles between the the cell periphery or within the perinuclear area (Figure 5c, d).

**Figure 5.**
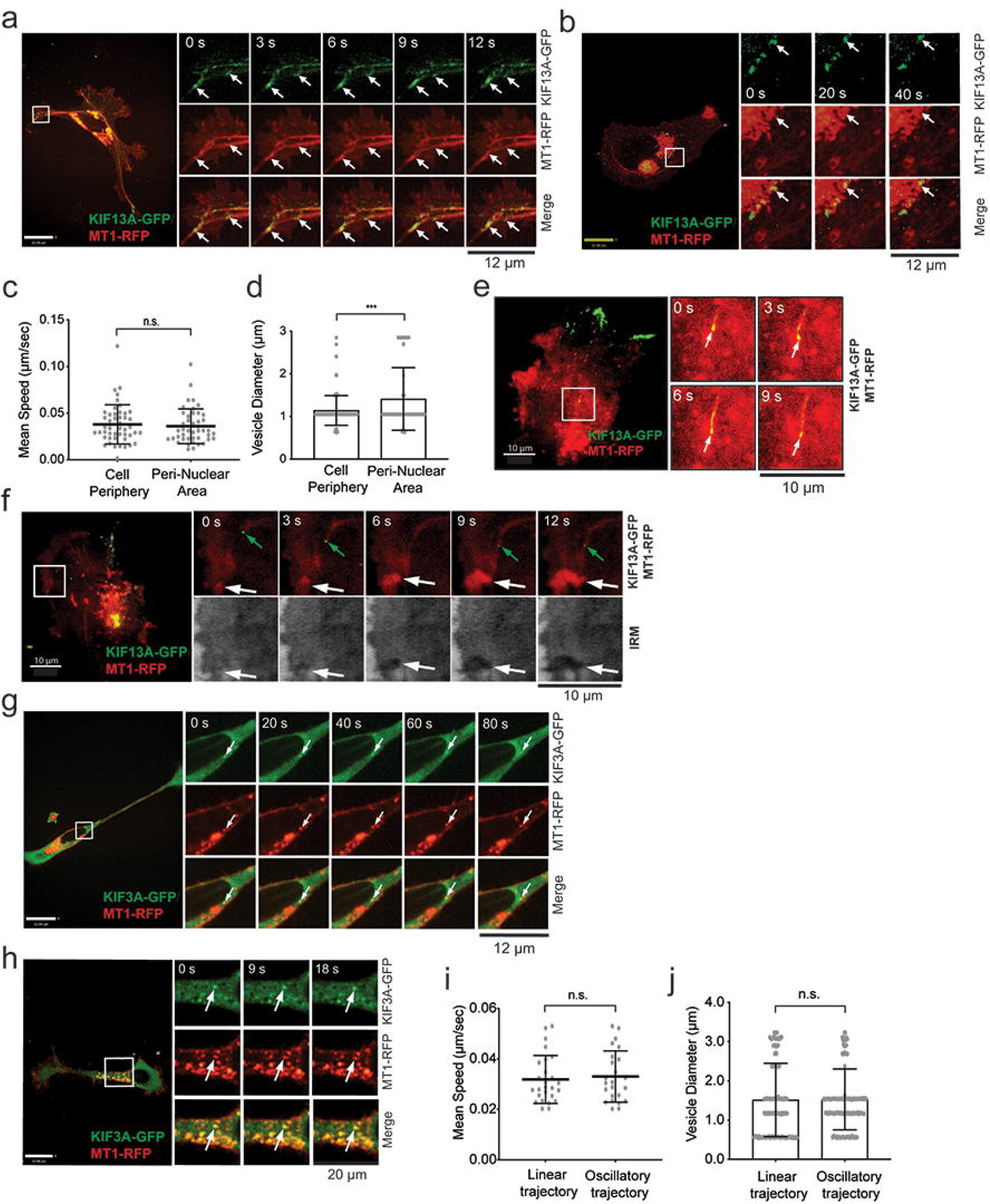
MT1-RFP was co-trafficked with KIF13A-GFP and KIF3A-GFP. **a, b.** HT-1080 cells were transfected with KIF13A-GFP (green) and MT1-RFP (red) and they were subjected to live cell imaging by confocal microscopy. Representative image time sequences are shown. White arrows point vesicles of interest. Scale bars are 22 μm. **c.** Quantification of mean velocity of vesicles of interest at the leading edge and nuclear area. Data are representative of five independent experiments and are calculated for 47 and 44 trajectories for the cell edge and the peri-nuclear area respectively. Unpaired T test with Welch’s correction. Data are shown as mean ± SD. n.s., non-significant. **d.** Quantification of vesicle diameter at the cell edge and peri-nuclear area. Data are representative of five independent experiments and are calculated for 129 vesicles for the leading edge and the peri-nuclear area. Data are shown as mean ± SD. *** P < 0.001. **e, f.** HT-1080 cells transfected with KIF13A-GFP (green) and MT1-RFP (red) were subjected to live cell imaging by TIRF microscopy. Representative image time sequences are shown. White arrows point vesicles of interest. IRM, internal reflection microscopy. Scale bar is 10 μm. **g, h.** HT-1080 cells transfected with KIF3A-GFP (green) and MT1-RFP (red) and they were subjected to live cell imaging by confocal microscopy. Representative image time sequences are shown. White arrows point vesicles of interest. Scale bars are 22 μm. **i.** Quantification of mean velocity of vesicles of interest with linear or oscillatory trajectories. Data are representative of five independent experiments and are calculated for 26 and 24 linear and oscillatory trajectories respectively. Unpaired T test with Welch’s correction. Data are shown as mean ± SD. n.s., non-significant. **j**. Quantification of vesicle diameter. Data are representative of five independent experiments and are calculated for 129 vesicles for each group. Unpaired T test with Welch’s correction was used. Data are shown as mean ± SD. n.s., non-significant.

We also carried out live cell imaging on TIRF microscopy (Figure 5e, *movie S3*) and the vesicles positive for KIF13A-GFP and MT1-RFP were detected at the cell-matrix interface. Combining time-lapse IRM with time-lapse TIRF microscopy, we also observed that MT1-RFP progressively accumulated at the growing adherent membrane on a gelatin-coated glass surface (Figure 5f). As discussed above, MT1-RFP-positive vesicles were again distributed in tubular-like structures, which pointed toward the growing highly adherent membrane (Figure 5f, *movie S4*)

Expression of KIF3A-GFP in HT-1080 cells did not affect the shape of MT1-RFP-positive vesicles (Figure 5g, h). The vesicles positive for KIF3A-GFP and MT1-RFP were either moving linearly (Figure 5g, *movie S5*) or oscillatory within the perinuclear region (Figure 5h, *movie S6*). Their mean velocities and diameters were equal, regardless of their trajectory (Figure 5i, j).

Vesicles double positive for KIF9-v1-GFP and MT1-RFP were detected within the perinuclear area of HT-1080 cells (Figure 6a). These vesicles did not move, but their fluorescence intensity faded over time, suggesting that the vesicles moved out of the focus plane, along the z-axis, most likely towards the dorsal side of the cell. As shown in Supplementary Figure 6b, no vesicles positive for both KIF9-v2-GFP and MT1-RFP could be detected. Likewise, we could not observe any MT1-RFP-positive vesicles trafficked by KIF1C-GFP (Figure 6c). Once more, KIF1C-GFP was accumulated at the trailing edge (Figure 6c).

**Figure 6.**
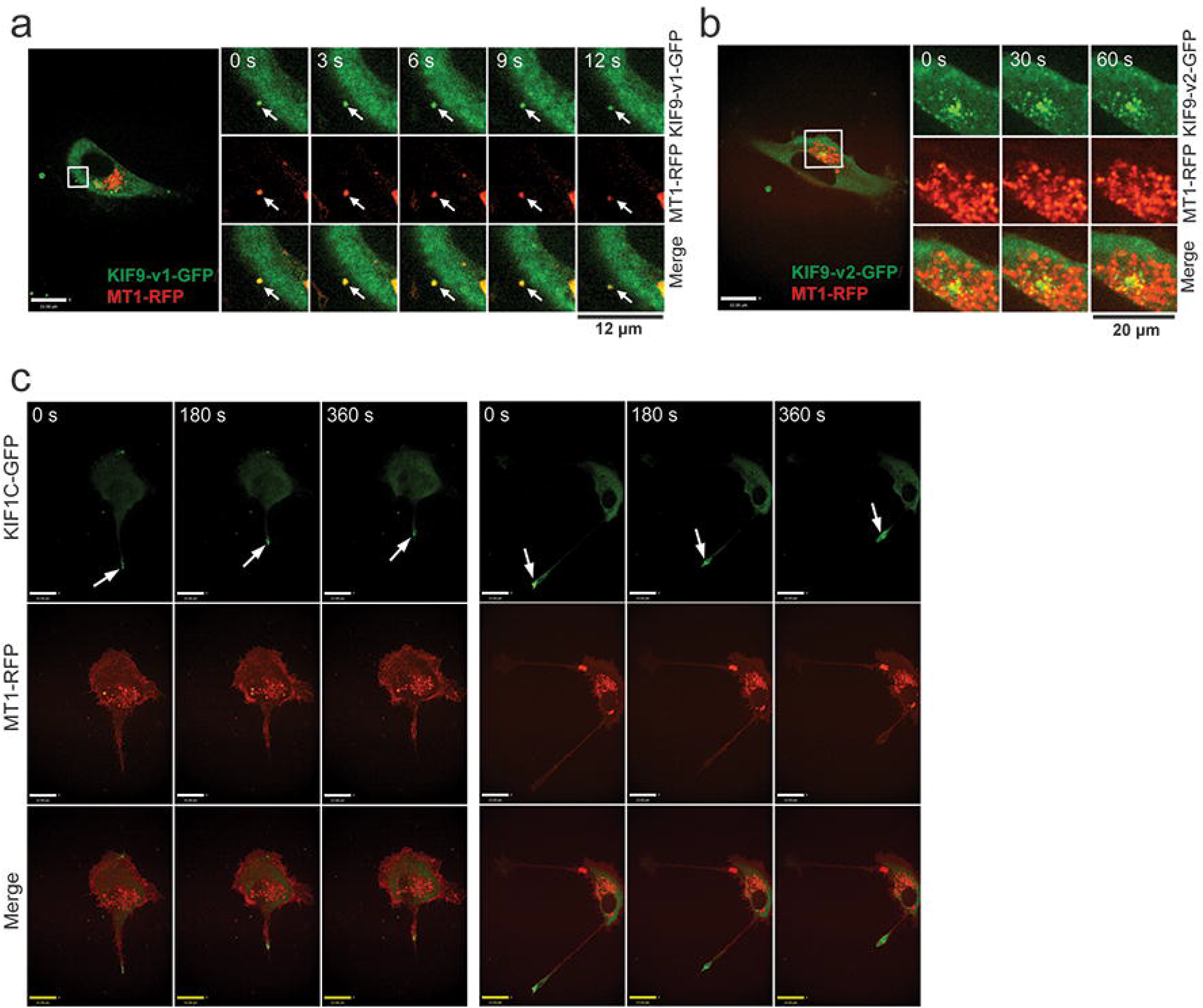
Time lapse imaging of cells expressing MT1-RFP, KIF9-v1-GFP, KIF9-v2-GFP, and KIF1C-GFP. **a.** HT-1080 cells transfected with KIF9-v1-GFP (green) and MT1-RFP (red) were subjected to live cell imaging by confocal microscopy. Representative image sequence is shown. White arrows point vesicles of interest. Scale bar is 22 μm. **b.** HT-1080 cells were transfected with KIF9-v2-GFP (green) and MT1-RFP (red) and they were subjected to live cell imaging by confocal microscopy. Representative image time sequence is shown. Scale bars are 22 μm. **c.** HT-1080 cells were transfected with KIF1C-GFP (green) and MT1-RFP (red) and they were subjected to live cell imaging by confocal microscopy. Representative image time sequences are shown. White arrows point trailing edges. Scale bars are 22 μm.

### Collaboration of KIF13A and KIF3A and competition of KIF9-v1 with KIF13A/KIF3A to transport MT1-MMP vesicles

We next investigated if KIF13A and KIF3A transport MT1-MMP vesicles independently or in collaboration. HT-1080 cells transfected for si-KIF13A, si-KIF3A, and a combination of the two were subjected to gelatin film degradation (Figure 7a, b, c). If these KIFs work independently, double knockdown of KIF3A and KIF13A should show additive inhibitory effect. However, if they work in collaboration, such an effect should not be observed. Upon KIF13A knockdown, gelatin film degradation was decreased by around 78%, while KIF3A knockdown by 49%. When the two knockdowns were combined, gelatin film degradation was decreased by 82%, which is the same level as KIF13A knockdown alone (Figure 7a, b). These data suggest that these two motor proteins function on the same pathway to deliver MT1-MMP to the cell surface.

**Figure 7.**
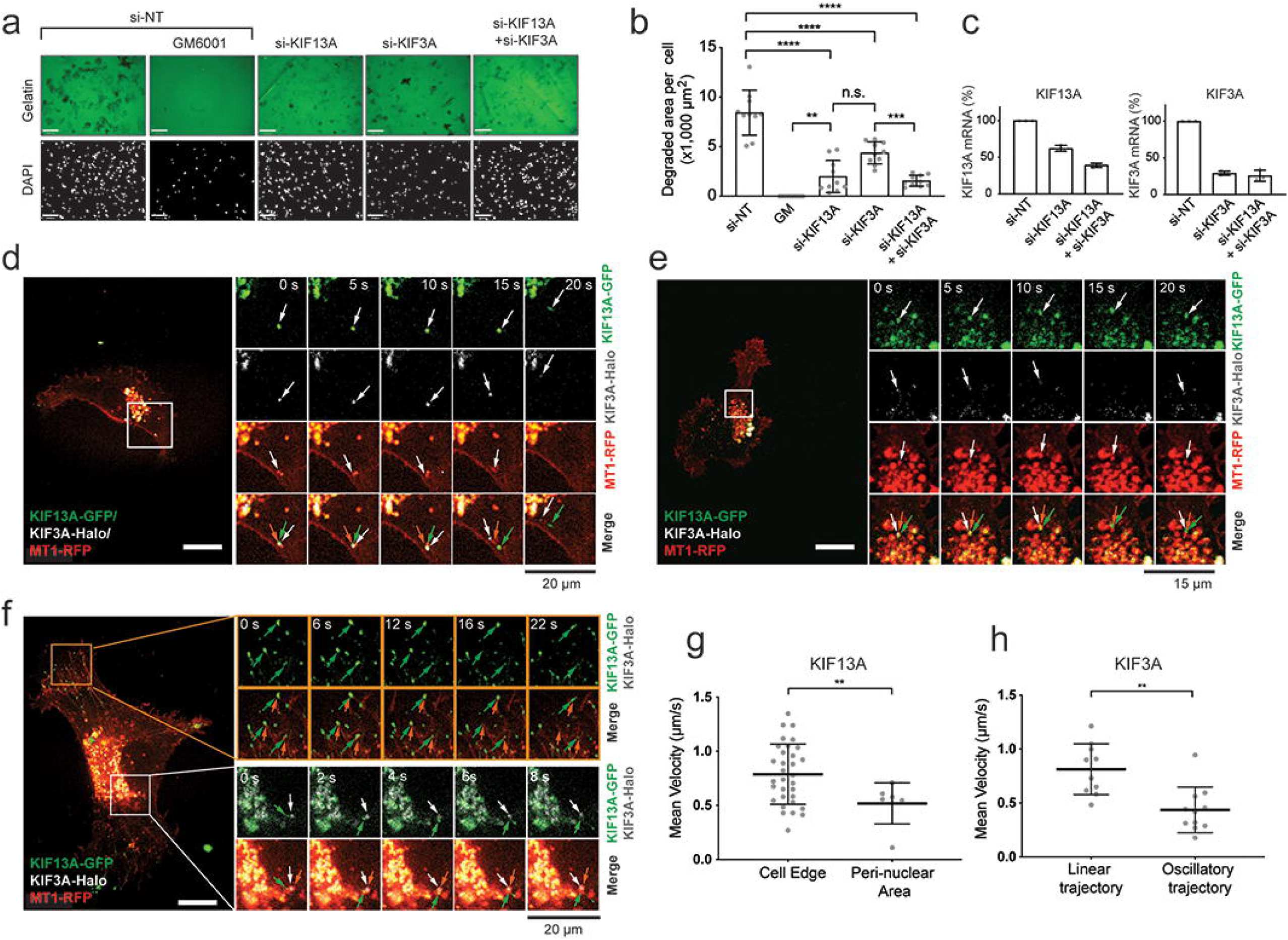
KIF13A and KIF3A transport MT1-MMP-containing vesicles. **a.** HT-1080 cells were transfected with siRNAs targeting the selected KIFs and subjected to gelatin film degradation assay in the presence or absence of GM6001 (10 μM). Scale bars are130 μm. **b**. Quantification of the degradation area (μm^2^) per cell in HT1080 cells transfected with the specified siRNAs. Data are presented as mean of 10 independent microscopic fields of view and are representative of three independent experiments. One-way ordinary Anova with Tuckey’s multiple comparisons test. Data are shown as mean ± SD. \*\*\*\**P* < 0.0001; \*\*\**P* < 0.001; \*\**P* < 0.01; n.s., non-significant. **c**. Efficiency of KIF knockdown was assessed by one-step RT-PCR. Quantification of KIF13A and KIF3A mRNA fold changes relative to GADPH mRNA. The data are representative of three independent experiments. **d. e, f.** HT-1080 cells were transfected with KIF13A-GFP (green), KIF3A-GFP (white) and MT1-RFP (red) and they were subjected to live cell imaging by confocal microscopy. Representative image time sequences are shown. Arrows point vesicles of interest. Scale bars are 22 μm. **g.** Quantification of mean velocities of MT1-RFP-containing vesicles transported by KIF13A-GFP at the cell edge and nuclear area. Data are representative of three independent experiments and are calculated for 32 and seven trajectories for the cell edge and the nuclear area respectively. Unpaired T test with Welch’s correction. Data are shown as mean ± SD. \*\**P* < 0.01; n.s., non-significant. **h.** Quantification of mean velocities of MT1-RFP-containing vesicles transported by KIF3A-GFP with linear or oscillatory 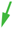trajectories. Data are representative of three independent experiments and are calculated for 10 linear and 11 oscillatory trajectories. Unpaired T test with Welch’s correction. Data are shown as mean ± SD. \*\**P* < 0.01, n.s., non-significant.

Next, HT-1080 cells expressing KIF13A-GFP, KIF3A-HaloTag, and MT1-RFP were subjected to time-lapse imaging. Vesicles positive for all three signals were detected within the perinuclear region (Figure 7d, e). As pointed by the arrows colored according to the different channels, these vesicles moved from the perinuclear area towards the cell periphery (Figure 7d). KIF13A-GFP and KIF3A-HaloTag co-localized with MT1-RFP-, containing vesicle until MT1-RFP fluorescence signal had faded, indicating that the MT1-RFP vesicle had fused with the plasma membrane (Figure 7d). Then, KIF13A-GFP and KIF3A-HaloTag independently trafficked back towards the center of the cell (Figure 7d). In other cases, these vesicles positive for KIF13A-GFP, KIF3A-HaloTag, and MT1-RFP were oscillating within the perinuclear region (Figure 7e). On the other hand, vesicles at the cell periphery were positive for only KIF13A-GFP and MT1-RFP (Figure 7f, *movie S7*). These vesicles were either moving within the cell edge or the perinuclear region. Interestingly, with KIF3A-HaloTag expression, KIF13A-GFP- and MT1-RFP-positive vesicles moved significantly faster at the cell edge within the perinuclear area (Figure 7g). With KIF13A-GFP expression, KIF3A-HaloTag- and MT1-RFP-positive vesicles moved significantly faster when they were proceeding with a linear trajectory than when they were oscillating (Figure 7h). Taken together, we concluded that KIF13A and KIF3A co-traffic MT1-MMP-containing vesicles around the perinuclear areas, and KIF13A takes over the vesicles at the periphery to traffic them towards the plasma membrane of the cell.

Given the role of KIF3A and KIF13A in MT1-MMP vesicle transport to degrade matrix, increased matrix-degrading activity upon KIF9 knockdown may be due to increased KIF3A and KIF13A-mediated vesicle transport of MT1-MMP. To test this hypothesis, *Kif9* was co-silenced with *Kif3a* or *Kif13a*. As shown in Figure 8, increased gelatin film degradation by KIF9 knockdown was significantly decreased upon co-silencing *Kif3a or Kif13a* genes. These data suggest that knockdown of KIF9 made MT1-MMP vesicles available for KIF3A- and KIF13A-dependent vesicle transport, resulting in increased matrix degradation.

**Figure 8.**
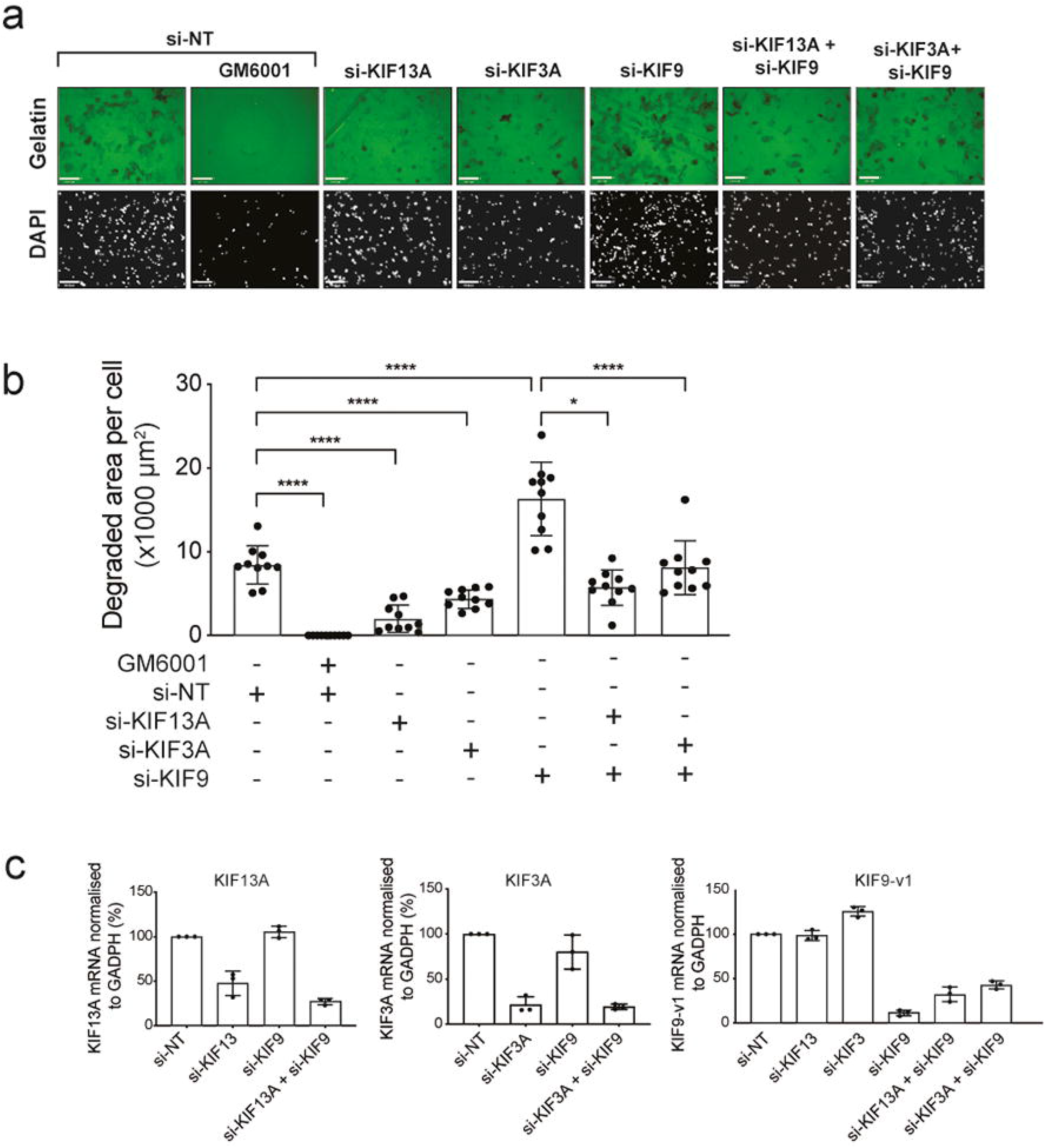
Increased gelatin film degradation upon KIF9 knockdown is due to increased KIF13A and KIF3A-dependent vesicle transport of MT1-MMP. **a**. HT1080 cells were transfected with siRNA for KIF13A, KIF3A and/or KIF9 and subjected to gelatin film degradation assay. Cells wer counterstained with DAPI (lower panels). Scale bar is 130μm. **b**. Quantification of the degradation area (μm^2^) per cell. Data are presented as the mean of ten independent microscopic fields of view and are representative of three independent experiments. *P* values were calculated by Student T test. Data are shown as mean ± S.D. \*\*\*\**p*>0.0001; \**p*>0.05. **c**. Efficiency of KIF knockdown by one-step RT-PCR. Quantification of KIF13A, KIF3A and KIF9-v1 mRNA fold changes relative to GADPH. The data are representative of three independent experiments.

### Role of KIF13A, KIF3A, KIF9 and KIF1C in MT1-MMP secretion in other cell types

We next investigated the role of these KIFs in other cell types. First, human rheumatoid arthritis synovial fibroblasts (RASFs) were examined since RASFs’ cell surface collagenolytic activity is solely MT1-MMP-dependent (Kaneko et al., 2016; Majkowska et al., 2017; Miller et al., 2009). RASFs from three different donors were transfected with siRNAs targeting these four KIFs and subjected to collagen film degradation assay. As shown in Figure 9 (a-c), knockdown of KIF13A and KIF3A significantly reduced collagen film degrading activity, while KIF9 knockdown enhanced it. KIF1C knockdown did not influence the collagen degradation. These data suggest that KIF13A, KIF3A, and KIF9 are similarly involved in MT1-MMP intracellular trafficking in RASFs.

**Figure 9.**
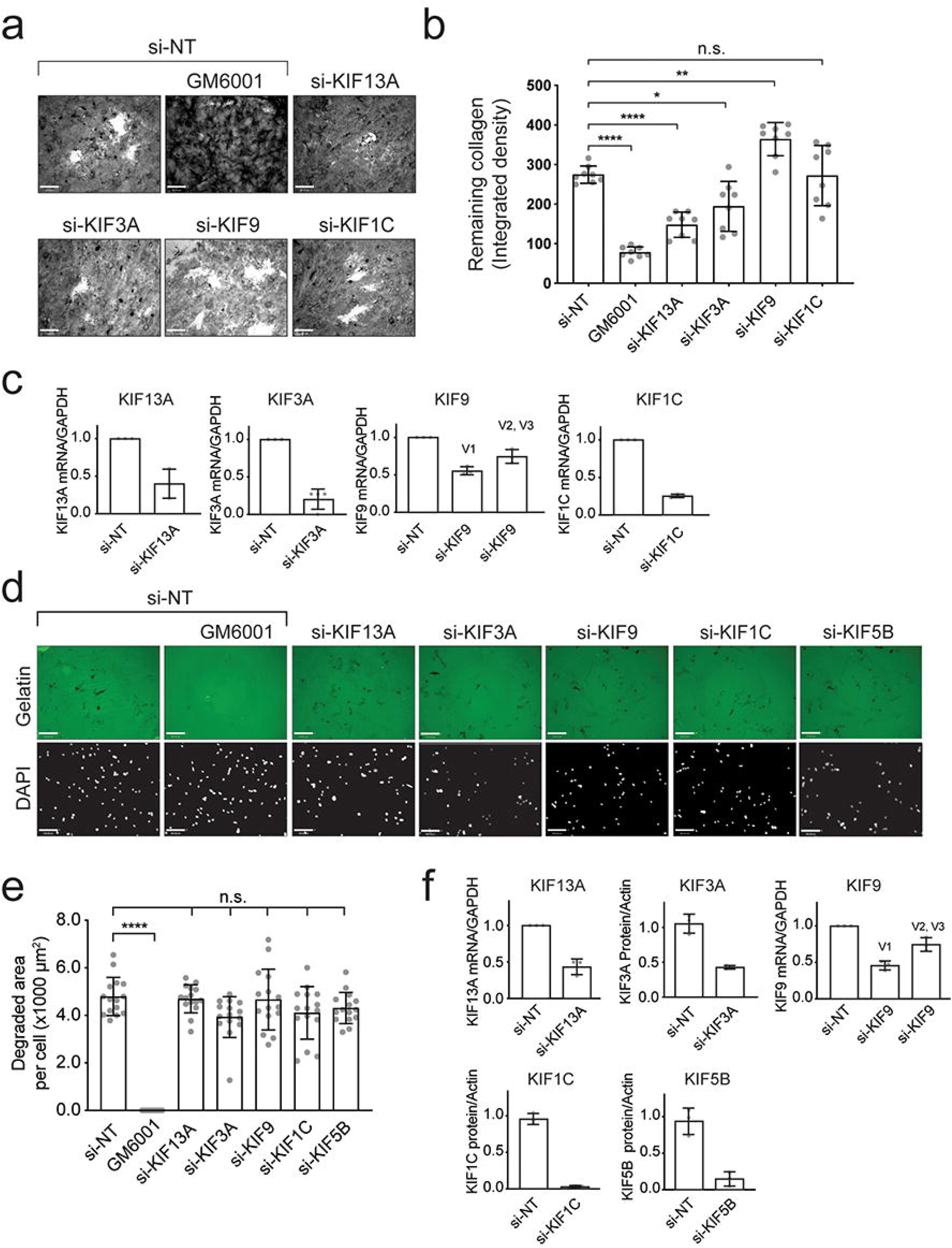
Effect of KIF knockdown on MT1-MMP activity in RASF and MDA-MB-231 cells. **a.** RASFs were transfected with siRNAs targeting the selected KIFs and subjected to collagen film degradation assay in the presence or absence of GM6001 (10 μM). Scale bars, 130 μm. **b.** Quantitative data of the integrated density of the collagen layer in RASFs transfected with the specified siRNAs. Data are presented as mean of eight independent microscopic fields of view and are representative of five independent experiments. One-way ordinary Anova with Tuckey’s multiple comparisons test. Data are shown as mean ± SD. \*\*\*\**P* < 0.0001, \*\**P* < 0.01, \**P* < 0.05, n.s., non-significant. **c.** Efficiency of KIF knockdown assessed by RT-PCR in RASFs were shown as graphs. The data are representative of three independent experiments. **d.** MDA-MB-231 cells were transfected with siRNAs targeting the indicated KIFs and subjected to gelatin film degradation assay in the presence or absence of GM6001 (10 μM). Scale bars are 130 μm. **e.** Quantitative data of the degradation area (μm^2^) per cell were shown as a graph. Data are presented as the mean of fifteen independent microscopic fields of view and are representative of five independent experiments. *P* values were calculated by One-way ordinary Anova with Tuckey’s multiple comparisons test. Data are shown as mean ± SD; \*\*\*\**P* < 0.0001. n.s., non-significant. **f.** Efficiency of knockdown for KIF13A and KIF9 were assessed by RT-PCR. The level of mRNAs relative to GADPH mRNA were shown as graphs. Efficiency of KIF3A and KIF1C knockdown were assessed by Western blot. Cell lysates were analysed by Western blotting. Quantification of the intensity of KIF3A, KIF1C and KIF5B protein bands relative to actin were shown as graphs. The data are representative of three independent experiments.

Next, triple-negative breast cancer cell line, MDA-MB-231 cells were examined. MDA-MB-231 cells have been characterized extensively for invadopodia-mediated cell invasion, which is also MT1-MMP-dependent (Castro-Castro et al., 2016; Ferrari et al., 2019; Frittoli et al., 2011; Infante et al., 2018; Poincloux et al., 2009). MDA-MB-231 cells were transfected with siRNAs targeting KIF13A, KIF3A, KIF9, KIF1C, and KIF5B and subjected to gelatin film degradation (Figure 9d-f). Interestingly, silencing these five *kinesin* genes, including KIF3A and KIF5B previously shown to associate with MT1-MMP-containing vesicles in MDA-MB-231 cells (Marchesin et al., 2015; Wang et al., 2017), did not impact MT1-MMP- and invadopodia-dependent gelatin film degradation (Figure 9d, e). These data suggest that roles of KIFs in trafficking MT1-MMP-containing vesicles are cell context-dependent.

### Genomic alterations of *Kif13a* and *Kif9* genes across The Cancer Genome Atlas (TCGA) studies

We next investigated if expression level of KIF3A, KIF13A and KIF9 have any correlation with cancer progression. Gene expression of KIF13A, KIF3A, and KIF9 across The Cancer Genome Atlas (TCGA) PanCancer Atlas studies were analyzed. Amplifications and deep deletions of *Kif13a, Kif3a*, and *Kif9* genes across the TCGA database were searched on cBioPortal (http://cbioportal.org)(Cerami et al., 2012; Gao et al., 2013). The term amplification indicates a high copy-number level per gene. Deep deletion implies a profound loss in the copy-number level per gene and often refers to a homozygous deletion. As shown in Figure 10a, the *Kif13a* gene was amplified in 1.3% of samples across the TCGA PanCancer Atlas studies and deleted in 0.2%. On the other hand, the *Kif9* gene was amplified in only 0.1% of samples and deleted in 0.5%. These gene alterations were distributed across several cancer types of TCGA (Figure 10b, c). More than seven in 100 women diagnosed with ovarian serous cystadenocarcinoma presented a *Kif13a* gene amplification (Figure 10b). *Kif13a* gene was also amplified in more than 6% and 4% of patients diagnosed with bladder urothelial cancer and diffuse large-B cell lymphoma (DLBC), respectively (Figure 10b). *Kif9* gene was deleted in over 6 out of 100 patients diagnosed with DLBC and in almost 3% of patients with kidney renal clear cell carcinoma (ccRCC) (Figure 9c). Next, we used the Pathology Atlas (https://www.proteinatlas.org/humanproteome/pathology)(Uhlen et al., 2017) to investigate whether KIF13A and KIF9 are considered prognostic markers for specific cancer types. We performed Kaplan-Meier survival analysis of liver cancer patients with low or high mRNA expression of the *Kif13a* gene through the Pathology Atlas. We found that patients with high expression of the *Kif13a* gene had a significantly lower survival rate (Figure 10d). We also performed the same analysis of renal and colorectal cancer patients with low or high mRNA expression of the *Kif9* gene. We found that patients with low expression had a significantly lower survival rate (Figure 10e, f).

**Figure 10.**
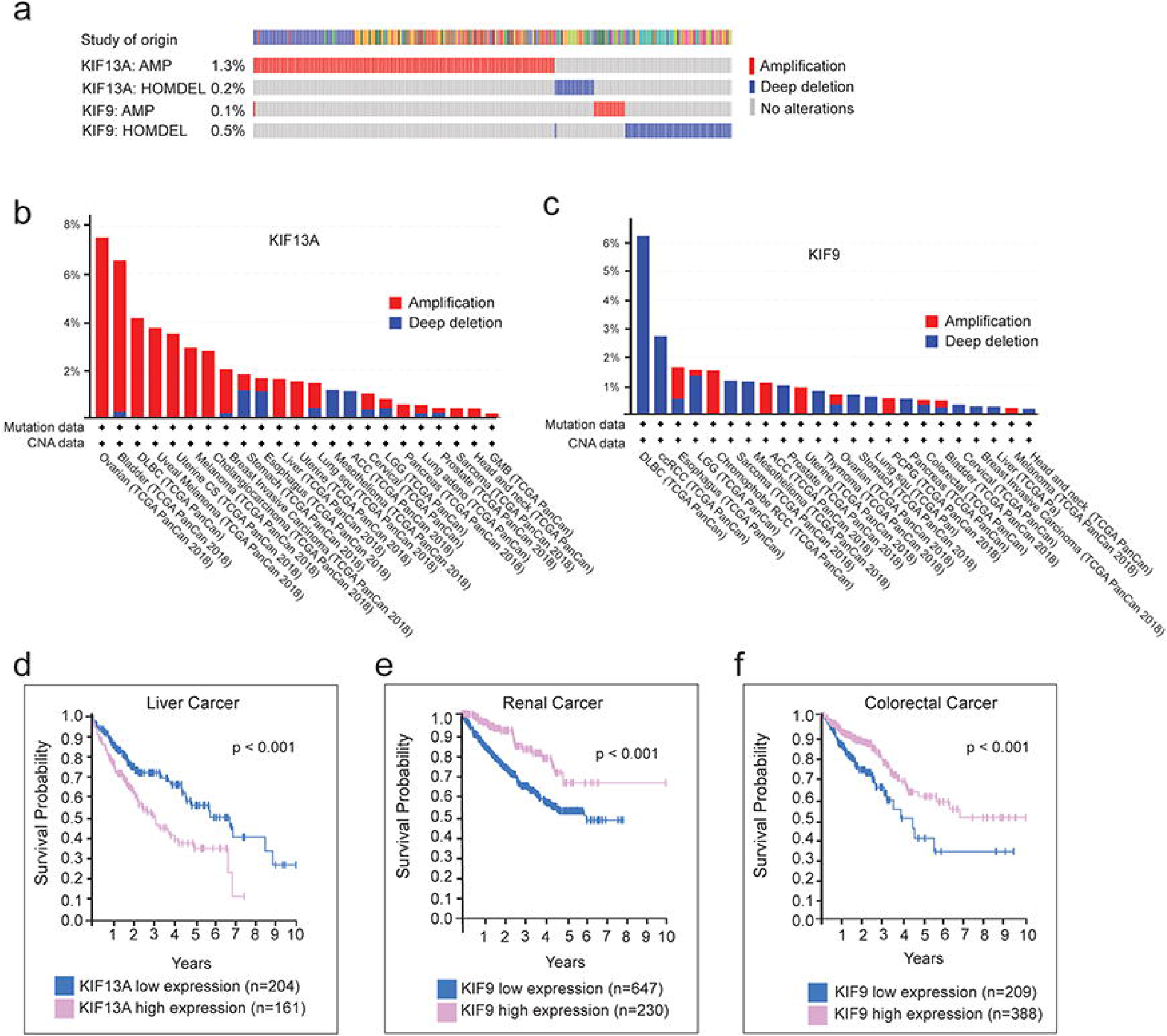
Genomic analysis of KIF13A and KIF9. **a.** The oncoprints of KIF13A and KIF9 were identified. Genetic alterations of KIF13A and KIF9. The columns represent patients from studies within TCGA and the rows gene alterations like amplification and deep deletion. **b.** Genetic alterations of KIF13A summarised according to the study type. Amplifications are shown in red and deep deletions are blue. **c.** Genetic alterations of KIF9 summarised according to the study type. Amplifications are shown in red and deep deletions are blue. **d.** Kaplan-Meier survival analysis of liver cancer patients with low or high mRNA expression of KIF13A. P value was obtained with log-rank test. **e.** Kaplan-Meier survival analysis of renal cancer patients with low or high mRNA expression of KIF9. *P* value was obtained with log-rank test. **f.** Kaplan-Meier survival analysis of colorectal cancer patients with low or high mRNA expression of KIF9. *P* value was obtained with log-rank test.

*Kif3a* gene was amplified in 0.5% of cases across the TCGA PanCancer Atlas studies and deleted in 0.3% of them (Figure S3a). 5% of patients with ccRCC presented an amplification of the *Kif3a* gene. However, we could not assess whether this gene alteration was associated with a change to the overall patients’ survival (Figure S3b). According to our search on the Pathology Atlas, *Kif3a* gene alterations are not considered prognostic markers for different cancer types.

## DISCUSSION

When invading cancer cells encounter a physical ECM barrier with narrow gaps of less than 7 μm in diameter, they utilize MT1-MMP to break through the barrier and create a path for migration (Wolf et al., 2013). In this process, cells directly secrete MT1-MMP to the invading edge by trafficking MT1-MMP-containing vesicles along microtubules. Here, we have identified three kinesin motor proteins, KIF3A, KIF13A, and KIF9, directly involved in the trafficking of MT1-MMP-containing vesicles to the leading edge.

Although knockdown of KIF1C significantly reduced gelatin and collagen film degradation like KIF3A and KIF13A knockdown, we could not get the evidence that KIF1C was directly involved in MT1-MMP vesicle transport. MT1-RFP and KIF1C-GFP did not co-localize when cells were cultured either on a gelatin film or within 3D collagen gel, and there was a strong tendency of KIF1C to localize at the trailing edge of migrating cells rather than at the leading edge. A similar conclusion was made by Weisner *et al.(Wiesner* et al., 2010), as they observed that KIF1C knockdown decreased MT1-MMP-dependent gelatin film degradation at podosomes, but KIF1C did not co-localize with MT1-MMP vesicles. KIF1C may affect MT1-MMP vesicle transport indirectly, by potentially influencing cell migration machinery, as knockdown of KIF1C, but not other KIFs, reduced cell migration on plastic significantly (Figure 3c), which is an MT1-MMP-independent process. Therefore, KIF3A and KIF13A are the major KIFs that directly traffic MT1-MMP vesicles to the leading edge in HT1080 cells.

Interestingly, the same effect of knockdown of KIF3A, KIF13A and KIF9 was also observed for proMMP-2 activation activity. We and others have previously reported that homodimer formation of MT1-MMP is essential for proMMP-2 activation on the cell surface to form a proMMP-2 activation complex of (MT1-MMP)_2_-TIMP-2-proMMP-2 (Gifford and Itoh, 2019; Itoh, 2015; Itoh et al., 2001; Lehti et al., 2002). We have previously reported that homodimerization of MT1-MMP occurs consistently at the leading edge of invading cells regulated through the re-organization of the cytoskeletal actin by Rac1 and Cdc42 (Itoh et al., 2011). Thus, the leading-edge localization of MT1-MMP via the vesicle transport by KIF3A and KIF13A may closely associate with proMMP-2 activation activity.

Our data suggest that MT1-MMP vesicles were initially trafficked by both KIF3A and KIF13A in coordination around the perinuclear area, and then KIF13A alone further transports the vesicles to the leading edge. To our knowledge, this is the first example of coordinated vesicle transport between KIF3A and KIF13A. KIF3A belongs to the kinesin-2 subfamily and has been shown to play a crucial role in the primary cilium formation (Hirokawa, 2000). It was shown that KIF3 is involved in post-Golgi transport of N-cadherin/*β*-catenin, which plays a role in the adhesion of neuronal progenitor cells (Teng et al., 2005). KIF13A belongs to the kinesin-3 subfamily and is known to mediate vesicle transport from the endosomal recycling compartment to the plasma membrane (Hirokawa et al., 2009; Nakagawa et al., 2000; Perez Bay et al., 2013). It has been shown that KIF13A, together with Rab22A and BLOCK1/2, plays a central role in the biogenesis of recycling endosomes (Shakya et al., 2018). Trans-Golgi network (TGN) is the last station of the secretory pathway of proteins, and after TGN, the cargo can be trafficked to three destinations: 1) the plasma membrane, 2) the secretory granules, and 3) the endosomes (Le Borgne and Hoflack, 1998). MT1-MMP is stored in endosomes upon endocytosis (Jiang et al., 2001; Planchon et al., 2018; Remacle et al., 2003), and during exocytosis (Pedersen et al., 2020). With our data that KIF13A knockdown significantly inhibited MT1-MMP activity on the cell surface, we speculate that KIF3A and KIF13A transport MT1-MMP vesicles from TGN to the endosome, and KIF13A further transports the vesicles from the endosomes to the plasma membrane.

There are three splicing variants of KIF9, KIF9-v1, -v2, and -v3. KIF9-v2 and -v3 encode identical amino acid sequences, while KIF9-v1 is 65 amino acids shorter in the coiled-coil region. Upon KIF9 knockdown using smart pool siRNA, KIF9-v1 was selectively knocked down. Interestingly co-localization with MT1-MMP vesicles was only observed with KIF9-v1, but not KIF9-v2. Therefore, increased gelatin and collagen film degradation and proMMP-2 activation resulted from KIF9 knockdown were most likely attributed to KIF9-v1 knockdown, which is the first example of functional difference between KIF9 variants. However, it is not clear why KIF9-v1, but not KIF9-v2, is selectively involved in MT1-MMP vesicle transport since KIF9-v1 and v2 share the same C-terminal tail region, where these KIFs are thought to interact with the cargo. Further investigation is required.

Live-cell imaging data indicated that the vesicles positive for MT1-RFP and KIF9-v1-GFP did not move much in a lateral direction. However, they faded out from the focus plane over time, suggesting that KIF9 may carry the vesicles to the dorsal side of the cell surface. Increased gelatin film degradation upon KIF9 knockdown was inhibited by co-silencing *kif13a* or *kif3a* genes, suggesting that the increased degradation is due to increased KIF13A- and KIF3A-dependent MT1-MMP vesicle transport to the matrix attachment sites. It is thus possible that KIF9-v1 competes with KIF13A and KIF3A to bring MT1-MMP-containing vesicles to different plasma membrane domains. Upon KIF9-v1 knockdown, vesicle transport by KIF13A and KIF3A starts to dominate, resulting in increased matrix degradation.

KIF9 knockdown increased cellular invasion in trans-well invasion assay, with the same tendency observed for gelatin and collagen film degradations. However, KIF9 knockdown significantly reduced cell migration in microcarrier beads invasion assay. This seemingly contradictory result can be due to differences between the two invasion assay systems. The first difference is whether cells face the collagen only at the ventral side (trans-well system) or surrounded by collagen fibrils (micro-carrier beads system). Another difference can be whether the migration is chemoattractant-guided or matrix-guided. In the trans-well system, the lower chamber contained 10% FBS, while the upper chamber was serum-free. Thus, cells in the upper chamber are attracted by the serum components such as lysophosphatidic acid (Idzko et al., 2004) in the lower chamber. In the microcarrier beads invasion assay, there are no chemoattractant gradients. Thus, cells are instead guided by collagen fibrils. When cells migrate through the 3D collagen matrix, cells enlarge the gaps of collagen fibrils by degrading it at the middle of pseudopods in a ring shape (Wolf et al., 2013; Wolf et al., 2007). KIF9 may be involved in the vesicle trafficking of MT1-MMP to these regions of the plasma membrane in coordination with KIF3A and KIF13A. KIF9 was previously shown to be involved in gelatin degradation at podosomes in macrophages (Cornfine et al., 2011). In that report, they found that reggie/flotillin interacts with the C-terminal cargo binding domain of KIF9, and knockdown of KIF9 or flotillin significantly reduced gelatin degradation at the podosomes (Cornfine et al., 2011). However, it was concluded that KIF9 was not directly involved in MT1-MMP vesicle transport to the podosomes in macrophages (Cornfine et al., 2011; Wiesner et al., 2010).

Our data in this study indicate that knockdown of KIF3A and KIF13A in human RASFs significantly reduced collagen film degradation. In contrast, KIF9 knockdown enhanced it (Figure 8), suggesting that RASFs share the same vesicle transport mechanism with HT1080 cells. On the other hand, knockdown of KIF3A, KIF13A, KIF9 did not influence invadopodia-mediated gelatin film degradation in MDA-MB231 cells (Figure 9d, e). Thus, motor proteins responsible for MT1-MMP vesicle transport are cell context-dependent. It has been reported that KIF5B and KIF3A/KIF3B are the responsible KIFs in trafficking MT1-MMP vesicles to the podosome in macrophages (Wiesner et al., 2010). KIF5B and KIF3A were also shown to mediate MT1-MMP vesicle transport to the invadopodia in MDA-MB231(Marchesin et al., 2015). However, our data indicate that knockdown of none of the KIFs we examined, including KIF5B, KIF13A, KIF3A, KIF9, and KIF1C, influenced invadopodia-mediated gelatin film degradation in MDA-MB231 cells. In HT1080 cells, KIF5B knockdown did not influence gelatin film degradation as well. One of the differences between these reports and our data is that we have analyzed endogenous MT1-MMP activity on the cell surface in both MDA-MB231 and HT1080 cells, while in these reports, KIF5B involvement was investigated on ectopically expressed mCherry-tagged MT1-MMP in both macrophages and MDA-MB231 (Marchesin et al., 2015; Wiesner et al., 2010). Since mCherry was inserted at the C-terminus of the cytoplasmic domain of MT1-MMP (Marchesin et al., 2015; Wiesner et al., 2010), the vesicle transport pathway of MT1-mCherry may differ from endogenous wild-type MT1-MMP. It is also possible that overexpressed enzyme may gain alternative vesicle trafficking pathways. Another possibility might be related to different cell types and culture conditions. Macrophages may potentially have a different mechanism of MT1-MMP vesicle trafficking from HT1080 cells, and MDA-MB231 cells in different lab may gain different mechanism. Nevertheless, we have concluded that KIF5B is not involved in vesicle transport of endogenous MT1-MMP in HT1080 cells.

The analysis of the TCGA database revealed that the expression profile of KIF13A and KIF9 is altered in several cancer types. Amplification is the most common *Kif13a* gene alteration detected across the TCGA database. KIF13A is reported by the Pathology Atlas (https://www.proteinatlas.org/humanproteome/pathology) (Uhlen et al., 2017) as an unfavorable prognostic marker for liver cancer patients. As shown in Figure 10, liver cancer patients with a high expression of KIF13A had a significantly lower survival probability than those with a low expression. Our data suggest that KIF13A is a crucial player of MT1-MMP intracellular trafficking pathways and mediate MT1-MMP-dependent invasion of cancer cells. Therefore, a pro-tumorigenic role of *Kif13a* maybe its involvement in MT1-MMP trafficking to the cell surface. KIF9 is mainly deleted rather than amplified across the TCGA database. According to the Pathology Atlas, it is a favorable prognostic marker in renal and colorectal cancers (Figure 10e and 10f). Thus, KIF9 may have an anti-tumorigenic potential for these two cancer types. These data highlight that KIF13A and KIF9 may have cancer type-specific roles, at least partly due to their roles in MT1-MMP intracellular trafficking. Chandrasekaran et al. (Chandrasekaran et al., 2015) used cBioPortal to search for kinesin gene alterations in TCGA. They reported that the *Kif13a* gene was amplified in more than 10% of patients diagnosed with serous ovarian adenocarcinoma or urothelial bladder carcinoma(Chandrasekaran et al., 2015). They also found that 12% of patients with clear renal carcinoma had a homozygous deletion of the *Kif9* gene(Chandrasekaran et al., 2015). These values were slightly different from our analysis of the TCGA database. The discrepancies are likely due to the large number of data added to the database after publication. Cho *et al*.(Cho et al., 2019) performed an integrated analysis of specific kinesin’s clinical significance, including KIF9, in low-grade glioma and glioblastoma. They showed that high KIF9 expression is linked to cancer progression and significantly lower survival probability, especially for patients diagnosed with glioblastoma(Cho et al., 2019). Thus, the role of KIF9 may be cancer-specific.

In conclusion, we have identified KIF3A, KIF13A, and KIF9-v1 as the major KIFs that traffic MT1-MMP vesicles in HT1080 cells. KIF3A and KIF13A collaborate and transport MT1-MMP to the leading edge, while KIF9-v1 seems to compete with KIF13A and KIF3A by trafficking MT1-MMP-containing vesicles to non-leading edge membrane structures. Our findings revealed novel mechanisms of interplay between different KIFs, which contribute to the understanding of vesicle transport mechanisms during cancer invasion.

## MATERIALS AND METHODS

### Plasmid constructs

MT1-RFP in pSG5 vector (Stratagene) was generated as described previously (Itoh et al., 2011). cDNAs encoding human KIF13A, KIF3A, KIF9-v1, KIF9-v2 and KIF1C were amplified by PCR using a cDNA library from HT-1080 cells as template. The AcGFP was inserted at the N-terminus of each KIFs with three glycine as linker. The mutants were constructed by the overlap extension PCR method. KIF13A-GFP (forward primer for AcGFP: 5’-TAGGAGCTCGGTACCGCCGCCACCATGGTGAGCAAGGGCG-3’; reverse primer: 5’-CCATTCCACCTCCCTTGTACAGCTATCCATGC-3’; forward chimera primer for AcGFP-KIF13A: 5’-TAGGAGCTCGGTACCGCCGCCACCATGGTGAGCGCAAGGGCG-3’; reverse flanking primer: 5’-TAGCCCGGGTCACTTGTACAGCTCATCCATGC-3’). KIF3A-GFP (forward flanking primer: 5’-ATACGACTCACTATAGGGCGAATTCGAGCCACCATGGTGAGCAAGGGCGCC-3’; reverse primer: 5’-CGGCATTCCACCTCCCTTGTACACTCATCCATGCCGTG-3’; forward chimera primer for AcGFP-KIF3A: 5’-TACAAGGGAGGTGGAATGCCGATCGGTAAATCAGA-3’; flanking reverse primer: 5’-CCTCTTCATCATCATCTTCC-3’), KIF9-v2 (forward flanking primer: 5’-AATTCGAGCTCGGTACCCAGATCTGCCACCATGGTGAGCAAGGGCG-3’; reverse primer: 5’-CCCATTCCACCTCCCTTGTACAGCTCATCCATGCCG-3’; forward chimera primer for AcGFP-KIF9-v2: 5’-ACAAGGGAGGTGGAATGGGTACTAGGAAAAAAGTTC-3’; flanking reverse primer: 5’-AATAAGATCTGGATCCCCCTATTTTCTATGTGCCTGCTG-3’) and KIF1C-GFP (forward flanking primer: 5’-AARRCGAGCTCGGTACCCGCCACCATGGTGAGCAAGGGCGCC-3’; reverse primer: 5’-GCCATTCCACCTCCCTTGTACAGCTCATCCATGCCGTG-3’; forward chimera primer for AcGFP-KIF1C: 5’-TACAAGGGAGGTGGAATGGCTGGTGCCTCGGTCAA-3’; flanking reverse primer: 5’-AATAAGATCTGGATCCCCTCACACAGCTGCCCCACTCTC-3’). KIF9-v1-GFP was generated by restriction enzyme cloning. KIF3A-HaloTag and KIF9-v2-HaloTag were generated by sub-cloning KIF3A and KIF9-v2 into pHTN HaloTag CMV-neo (Promega). KIF13A, KIF13A-GFP, KIF9-v1, KIF9-v2, KIF9-v1-GFP, KIF9-v2-GFP, KIF1C, and KIF1C-GFP were subcloned into pSG5 vector.

### Cell culture, transient transfection and siRNA treatment

HT-1080 human fibrosarcoma cells (ECACC) were cultured in Dulbecco’s modified Eagle’s medium (DMEM) (Lonza), containing 10% FBS (Gibco), penicillin/ streptomycin (P/S) (PAA). Rheumatoid arthritis synovial fibroblasts derived from three different patients were cultured in DMEM supplemented with 20% FBS and P/S (Miller et al., 2009). HT-1080 cells were transfected with plasmid constructs using Trans-IT2020 (Mirus Bio) according to the manufacturer’s instructions. Gene silencing was performed by transfection of SMARTpool ON-TARGETplus siRNA (Dharmacon, Thermo Fisher, Waltham, US) using INTERFERin (Polyplus-transfection) according to the manufacturer’s instructions. Non-targeting siRNA (NT siRNA) was also purchased from Dharmacon. Gene silencing effectiveness was tested by Western blotting (WB) or RT-PCR. Cells were subjected for the experiments after 72 hours of transfection.

### SDS-PAGE and Western blotting

Cell lysates were prepared by directly dissolving in 1xSDS loading buffer containing 0.1% 2-mercaptoethanol. Cell lysates were subjected to SDS-PAGE and the proteins in the gel were transferred to a PBDF membrane using Trans-Blot Turbo Transfer System (Bio-Rad). After probing the membrane with the primary antibodies, the bands were visualised using fluorescently-labelled secondary antibodies (LI-COR). Membranes were scanned by Odyssey CLx imaging system (LI-COR). The band intensities were quantified by ImageJ software (National Institutes of Health). Actin or tubulin bands were used for normalisation. Mean band intensities were plotted using Prism (GraphPad software, Inc.) and statistical significance was calculated using either a parametric unpaired T test or a one-way ordinary ANOVA with Tuckey’s multiple comparison test. The following primary antibodies were used: rabbit anti-MT1-MMP monoclonal antibody (clone EP1264Y, ab51074, abcam), mouse anti-MT1-MMP monoclonal antibody (clone 222-1D8), rabbit anti-KIF1C polyclonal antibody (ab72238, abcam), rabbit anti-KIF3A polyclonal antibody (ab11259, abcam), mouse anti-actin monoclonal antibody (clone C4, sc47778, Santa Cruz), mouse anti-tubuline antibody (clone B-7, sc5286, Santa Crutz). The following secondary antibodies were used: IRDye 680RD goat anti-mouse IgG (H + L) (926-68070, LI-COR), IRDye 800CW goat ant-rabbit IgG (H + L) (926-32211, LI-COR).

### RT-PCR

Total RNA was isolated using the RNAqueous Micro kit (Invitrogen) according to the manufacturer’s instructions. RNA was reversed-transcribed using the High Capacity cDNA Reverse Transcription kit from AB applied Biosciences (Thermo Fisher). Resulted cDNA was used as a template for PCR using the Dream Taq Polymerase (Thermo Fisher). GADPH was used as housekeeping gene. The following primers were used: KIF13A (forward: 5’-TTTCCAGTAGGAGGAGTC-3’; reverse: 5’-AAGTTGTTGCGGTGAAGG-3’), KIF3A (forward: 5’-TGCAAAGTCAGAGATGGC-3’; reverse: AGCTGCCATTCTCCTATG-3’), KIF9 (forward: 5’-CCCGGACCTTATCAGAGGAAAAG-3’; reverse (v1): 5’-GGTGTCGGGCCTGAGTGG-3’; reverse (v2): 5’-GGATGGGACAAGCTGGGTC-3’), KIF1C (forward: 5’-TTCCAGCCCAAAAAGCAC-3’; reverse: 5’-CGGACCTTCTCTCTCATC-3’), KIF5B (forward: 5’-GCTACAAGAGTTAAAAAGAGTGCT-3’; reverse: 5’-TCACACTTGTTTGCCTCCTCCAG-3’), MT1-MMP (forward: 5’-GGGACCTACGTACCCACACA; reverse: 5’-TAGCGCTTCCTTCGAACATT-3’) AND GADPH (forward: 5’-TTCACCACCATGGAGAAGGC-3’; reverse: 5’-GGTCCCTCCGATGCCTGC-3’). RT-PCR gels were quantified by Fiji using the same protocol described above for the quantification of WB gels. GADPH bands were used for normalisation.

### Gelatin film degradation assay

Fluorescently-labelled gelatin coated coverslips were prepared as described previously (Evans and Itoh, 2007; Itoh et al., 2006). Cells were cultured atop of the gelatin-coated coverslips for 2-15 hours, depending on the assay, in the presence or absence of GM6001 (10 μM) (Elastin Products Company), TIMP-1 (200 nM), or DX-2400 (200 nM). TIMP-1 was a gift from Prof Gillian Murphy (University of Cambridge), and DX-2400 was a gift from Dyax Corp. Cells were then fixed in 4% formaldehyde (Sigma Aldrich) in PBS for 15 minutes and stained with DAPI (Sigma-Aldrich). Images were acquired with a Nikon microscope with using the 10X dry lens (NA = 0.3) on a Nikon TE2000-E microscope equipped with an ORCA-ER CCD camera (Hammamatsu Photonics) operated by Volocity Acquisition module software (Improvision). Gelatin film degradation was quantified using Fiji. Area of degradation was calculated using the gelatin fluorescence image which was converted to greyscale and thresholded. The threshold was set the same for all the pictures analysed, in order to have an objective mean of analysis. Area and area fraction were measured. The DAPI-stained corresponding image was converted to a binary image in order to count the number of cells in the microscopic field. The degraded area per cell was calculated as follow:

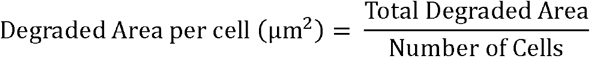

Mean degraded areas were plotted using Prism and statistical significance was calculated using one-way ordinary ANOVA with Tuckey’s multiple comparison test.

### Collagen film degradation assay

Collagen film degradation was carried out as described previously (Evans and Itoh, 2007; Itoh et al., 2006). Briefly, PureCol (bovine collagen type-I, Advanced Biomatrix) and Cellmatrix type I-A collagen (Nitta Gelatin) were mixed in the ratio 1:1. This mixture was neutralised and diluted to a 2 mg/ml. 12-well multi well plates were coated with 100 μl collagen solution/well and set for gelation. Cells were cultured on the collagen film for 72 hours in the presence or absence of GM6001 (10 μM), TIMP-1 (200 nM), or DX-2400 (200 nM). Cells were removed extensively by trypsin/EDTA (Lonza) and the remaining collagen layer was fixed with 4% formaldehyde in PBS and stained with R-250 Coomassie Brilliant Blue (Thermo Fisher). Representative images were acquired using the 10X dry lens (NA = 0.3) on the Nikon TE2000-E microscope equipped with an ORCA-ER CCD camera (Hammamatsu Photonics) operated by Volocity Acquisition module software (Improvision, PerkinElmer). Collagen degradation was quantified using Fiji by measuring the integrated density of the collagen layer. Mean results were plotted using Prism and statistical significance was calculated using one-way ordinary ANOVA with Tuckey’s multiple comparison test.

### Zymography

Gelatin zymography was conducted as reported previously (Evans and Itoh, 2007; Itoh et al., 2006). Enzyme activity was visualised directly on the gels as negative staining bands with Coomassie Blue. Pro-MMP-2 (P), intermediate MMP-2 (I) and active MMP-2 (A) bands were quantified using Fiji. The percentage of processed pro-MMP-2 over the total was calculated as follow:

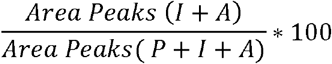

### Cell Surface biotinylation

Surface biotinylation was carried out using Sulfo-NHS-biotin (Thermo Fisher) as described previously(Itoh et al., 2001). Cells were cultured to confluency, and cell surface proteins were labelled with sulfo-NHS-biotin (Thermo-Fischer), followed by affinity precipitation of biotinylated molecules by streptavidin-conjugated Dyna beads (Thermo Fisher). The eluted samples were subjected to Western Blotting analyses using rabbit monoclonal anti-MT1-MMP antibody.

### Indirect immunofluorescent staining

Cells cultured atop of un-labelled or fluorescently labelled gelatin were fixed with 4% (v/v) formaldehyde in PBS for 5 min and blocked with 3% (w/v) BSA, 5% (v/v) goat serum in TBS (blocking solution) for 1 hour at room temperature (RT). Cells were then incubated with primary antibodies in blocking solution. The cells were further probed by Alexa488-conjugated goat anti-mouse IgG (Molecular Probes), DyLight650-conjugated goat anti-rabbit IgG (Thermo Scientific), Alexa568-conjugated Phalloidin (Molecular Probes, Thermo Fisher) or DAPI. The following primary antibodies were used for staining: rabbit anti-MT1-MMP monoclonal antibody (clone EP1264Y, abcam), mouse anti-human β1 integrin monoclonal antibody (clone 12G10, Millipore).

### Trans-well invasion assay

Trans-well invasion assay was performed as previously described(Palmisano and Itoh, 2010). Briefly, a 24-well insert with an 8-μm-pore membrane Trans-wells (VWR International Ltd) was coated with 50 μl of CellMatrix/PureCol collagen mixture (1:1, 2 mg/ml), incubated at 37°C for 1 hour to set the collagen and dried overnight at RT. Cells (2 × 10^4^/well) were seeded in the upper chamber and further cultured for 18 hours. Lower chamber media contained 10% FBS while upper chamber media was serum free. Invaded cells were stained with DAPI, imaged with fluorescence microscopy, and analyzed by Volocity software.

### Microcarrier beads invasion assay

Microcarrier beads invasion assay was carried out as described previously (Palmisano and Itoh, 2010). Cells were attached to gelatin-coated Cytodex 3 microcarrier beads (VWR) by preparing a cell/bead suspension which was incubated on a shaker for 6 hours at 37°C. Beads coated with cells were suspended in neutralised Cellmatrix collagen (final concentration 2 mg/ml) and incubated overnight in the presence or absence of GM6001 (10 μM). Invasion was analysed as the distance between a cell nucleus and the surface of the bead using the line tool of ImageJ. Migrated distances were calculated for fifty cells per treatment. Mean migrated distances were plotted using Prism and statistical significance was calculated using one-way ordinary ANOVA with Tuckey’s multiple comparison test.

### Wound-healing assay

Ibidi Culture-Inserts (Ibidi) consisting of two reservoirs separated by a 500-μm-width wall was placed in each well of a 24-well plate and six inserts were used for each treatment. Cells were seeded in the two reservoirs of the inserts and incubated until confluence. After the inserts were gently removed, cells were further cultured for 6 hours. Pictures were taken immediately after the inserts were removed and at the 6-hour time point. To measure the percentage of wound closure, ImageJ was employed. For each condition, the ROI corresponding to the initial wound-gap (ROI-I) and the one, delimited by the migration front after 6 hours of incubation (ROI-F), were calculated. The percentage of wound closure was calculated as follow:

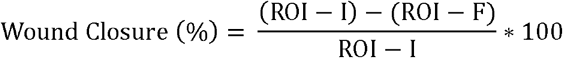

Mean percentages were plotted using Prism and statistical significance was calculated using one-way ordinary ANOVA with Tuckey’s multiple comparison test.

### Live cell imaging

Cells were transfected with GFP-tagged KIFs, Halo-tagged KIFs and MT1-RFP. After 36 hours they were seeded at a density of 4 × 10^3^/well on gelatin-coated glass-bottomed 8-well chambers (Ibidi). After 12 hours, live images were acquired by either confocal laser scanning microscopy or total internal reflection fluorescence (TIRF) microscopy (details specified below). Both microscopes were equipped with an environmental chamber to maintain temperature at 37°C and CO_2_ at 5%. Time-lapse images were acquired every 3 to 30 seconds, depending on the experiment.

### Image acquisition

All widefield images were captured on an inverted Nikon TE2000-E widefield microscope with Volocity Acquisition software (PerkinElmer). These objective lenses were used: 4X objective lens (Plan Fluor 4X/NA 0.13), 10X objective lens (UPLSAPO 10X/NA 0.30 DIC), and 20X objective lens (UPLSAPO 20X/NA 0.45). Confocal laser scanning microscopy imaging was performed on a PerkinElmer Spinning Disk Confocal Microscope based on a Nikon TE 2000-U Eclipse motorized inverted microscope with DIC optics. A 60X objective lens was used: 60X (Plan Apo 60×/NA 1.40). Volocity software (PerkinElmer, Coventry, UK) was used for Acquisition. For TIRFM an Olympus microscope in TIRF mode (CellTIRF-4Line system; Olympus), equipped with an EMCCD camera (Evolve) was used. A 150X objective lens was used (UPLSAPO 150 X/ NA 1.45).

### Data analysis

To measure the amount of either β1 integrin and MT1-MMP at the substrate-attachment side, the corresponding fluorescence intensities (FIs) were measured by ImageJ. TIRF images were analysed by defining a region of interest (ROI), corresponding to the cell body (ROI-CB), and a rectangular ROI in the background (ROI-B). FIs were calculated as follow:

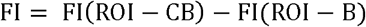

30 cells were analysed for each condition. ROI-B dimension and position was kept the same throughout the course of the analysis. Mean FIs were plotted using Prism and statistical significance was calculated using one-way ordinary ANOVA with Tuckey’s multiple comparison test.

To measure the overall amount of MT1-MMP on the cell surface of HT-1080 cells, FIs were analysed by Volocity measurement module software. Extended focus images of HT-1080 cells stained for cell surface endogenous MT1-MMP, captured by spinning disk confocal microscope were used. Thirty cells were analysed for each condition. Mean FIs were plotted using Prism and statistical significance was calculated using one-way ordinary ANOVA with Tuckey’s multiple comparison test.

Co-localisation between GFP-tagged KIFs and MT1-RFP was quantified using the Coloc 2 plugin in Fiji, which calculates Pearson’s correlation coefficients (PCCs). This procedure was used on two-colour channel images of cells cultured on 2D matrices and on two-colour channel stacks of cells cultured in 3D collagen matrices. Mean PCCs were plotted using Prism and statistical significance was calculated using one-way ordinary ANOVA with Tuckey’s multiple comparison test.

Tracking of MT1-RFP-containing vesicles trafficked by GFP-tagged KIFs was performed using the Track Mate plugin (Tinevez et al., 2017) in Fiji. Vesicles were identified using the LoG detector and their diameter was estimated at 1.5 μm. Tracking was performed by Simple LAP Tracker using 2 μm as linking and gap-closing maximum distances. The gap-closing maximum frame gap was set at 2. Mean velocities and vesicle diameters were calculated automatically by the plugin. Data were plotted using Prism and statistical significance was calculated by a parametric unpaired T test or a one-way ordinary ANOVA with Tuckey’s multiple comparison test.

To analyse genomic alterations of Kif13a, Kif3a and Kif9 genes in cancer, we employed cBioPortal (Cerami et al., 2012; Gao et al., 2013) (https://www.cbioportal.org). The Cancer Genome Atlas (TCGA) PanCancer Atlas studies were selected to visualise and analyse Kif13a, Kif3a and Kif9 gene alterations. The Human Protein Atlas (https://www.proteinatlas.org) was used to check whether Kif13a Kif3a or Kif9 genes were prognostic markers for specific cancer types.

## Supporting information

Subblemental table S1

## ACKNOWLEDGEMENT

We thank Prof Gillian Murphy for providing recombinant human TIMP-1. We thank Dyax Corp for providing us DX-2400. This work was funded by Cancer Research UK (Ref C1507/A12015) and DPhil studentship from Kennedy Trust of Rheumatology Research (KTRR). The Kennedy Institute of Rheumatology Cell Dynamics Platform (M. L. D. and Y. I.) was supported by KTRR and Wellcome 100262Z/12/Z (for the TIRF microscope). S. B. and M. L. D. were supported by ERC-2014-AdG 670930.

