## Supplementary material for "Coordination of KIF3A and KIF13A regulates leading edge localization of MT1-MMP to promote cancer cell invasion": Subblemental table S1

**Table S1. Selection of KIFs for screening**

There are 45 KIFs genes in human. Based on the literatures, 28 KIFs including splicing variant were excluded from the screening. The list of excluded KIFs and the reason with the references were shown.

| Excluded KIFs | | |
|---|---|---|
| Kinesin # | KIF # | Reason for being excluded with references |
| Kinesin-1 | KIF5A<br>KIF5C | Neuron specific(Kanai et al., 2000) |
| Kinesin-2 | KIF17 | Neuron specific(Guillaud et al., 2003) (Setou et al., 2000) |
| Kinesin-3 | KIF1A<br>KIF14<br>KIF16A<br>KIF16B<br>KIF28 | Neuron specific(Okada et al., 1995); Mitotic kinesins (Gruneberg et al., 2006); or function unknown |
| Kinesin-4 | KIF4A<br>KIF4B | Chromokinesins, expressed mostly in the juvenile brain (Kurasawa et al., 2004; Sekine et al., 1994). |
| Kinesin-5 | KIF11 | Mitotic kinesin(Kapitein et al., 2005; Sawin et al., 1992) |
| Kinesin-6 | KIF20A<br>KIF20B<br>KIF23 (v1,v2) | Mitotic kinesin (Fontijn et al., 2001). |
| Kinesin-7 | KIF10 | Mitotic kinesin(Ma et al., 2006) |
| Kinesin-8 | KIF19A<br>KIF19B<br>KIF18A<br>KIF18B | Unknown function(Miki et al., 2001; Stumpff et al., 2008); Mitotic kinesin(Mayr et al., 2007). |
| Kinesin-9 | KIF6 | Testis-specific(Nakagawa et al., 1997) |
| Kinesin-10 | KIF22 | Chromokinesin(Ohsugi et al., 2008) |
| Kinesin-11 | KIF26A<br>KIF26B | Microtubule-independent(Lillie and Brown, 1998) |
| Kinesin-12 | KIF12 | Mitotic kinesin(Lakshmikanth et al., 2004). |
| Kinesin-13 | KIF2A<br>KIF2B<br>KIF2C | M-type kinesin with microtubule-depolymerising activity (Homma et al., 2003) |
| Kinesin-14 | KIFC1<br>KIFC2<br>KIFC3<br>KIF25 (v1,v2) | C-type kinesin which moves towards the minus end of microtubule; implicated in acrosome formation (Yang and Sperry, 2003) (Yang et al., 2006). |
