## Supplemental Figs 1-3 for "Coordination of KIF3A and KIF13A regulates leading edge localization of MT1-MMP to promote cancer cell invasion"

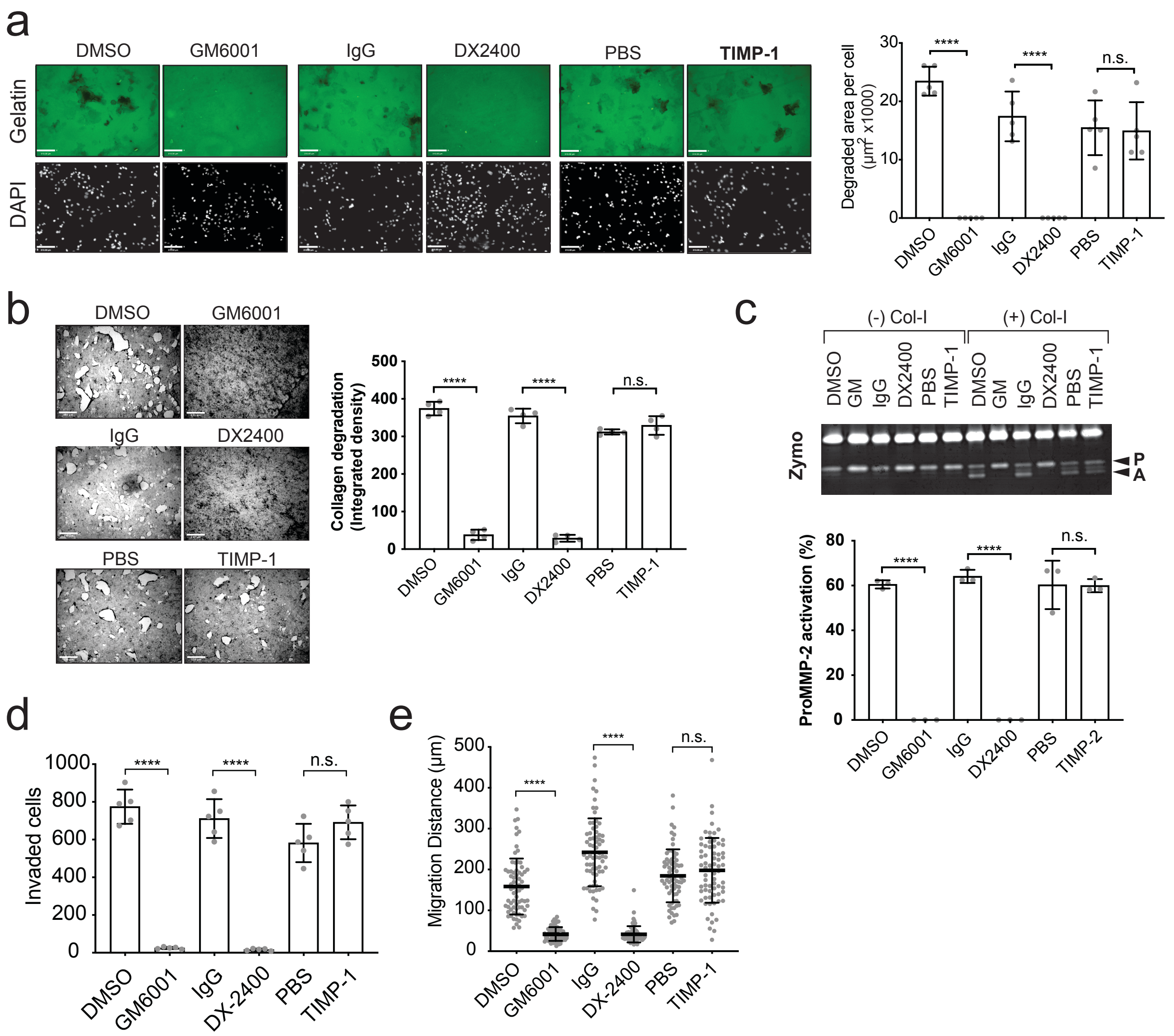


**Figure S1. Gelatin and collagen film degradation and proMMP-2 activation, cellular invasion measured by trans-well and microcarrier beads invasion assays are attributed to endogenous MT1-MMP in HT1080 cells.**

**a.** HT1080 cells were subjected to gelatin film degradation assay in the presence of GM6001 (10 μM), DX2400 (200 nM), TIMP-1 (200 nM) and the each control (DMSO, IgG, and PBS) (left panel). Scale bars, 130 μm. Quantitative data of the degradation area (μm2) per cell in HT1080 cells were shown as a graph (right panel). Data are presented as mean ± SD of five different field of the images (n=5). Data are representative of three independent experiments. One-way ordinary Anova with Tuckey's multiple comparisons test. **** P < 0.0001. n.s., non-significant.

**b.** HT1080 cells were subjected to collagen film degradation assay in the presence of GM6001 (10 μM), DX2400 (200 nM), TIMP-1 (200 nM) and the relative controls (right panel). Scale bars, 300 μm. Quantitative data of the integrated density of the collagen layer in HT1080 cells were shown as a gragh (right panel). Data are shown as mean ± SD (n=4) and are representative of three indipendent experiments. One-way ordinary Anova with Tuckey's multiple comparisons test. **** $P < 0.0001$, n.s., non significant.

**c.** HT1080 cells were cultivated in the presence or absence of collagen I (100 µg/ ml) with the above inhibitors for 24 h. Culture media from the assay were analysed by zymography (Zymo) (top panel). P, pro-MMP-2; A, active MMP-2. Quantification of the percentage of MMP-2 processed forms (active and intermediate forms) over the total MMP-2 (proforms, intermediate, and active forms) upon collagen stimulation are shown as a graph (bottom panel). Data are presented as mean ± SD (n=3) and the data shown is a representative of three indipendent experiments. One-way ordinary Anova with Tuckey's multiple comparisons test. **** $P < 0.0001$, n.s,. non significant.

**d.** HT1080 cells were subjected to transwell invasion assay in the presence of GM6001 (10 µM), DX2400 (200 nM), TIMP-1 (200nM) and relative control solutions. MT1-MMP-dependent cell invasion was quantified and shown as a graph. Data are presented as mean ± SD (n=6). Data are representative of three independent experiments.

**e.** HT1080 cells were subjected to micro carrier beads invasion assay in the presence of GM6001 (10 µM), DX2400 (200 nM), TIMP-1 (200nM) and each control. Distance of migration from the bead surface was quantified as described in the Method section and shown as a graph. Data are presented as a mean ± SD (n=100).

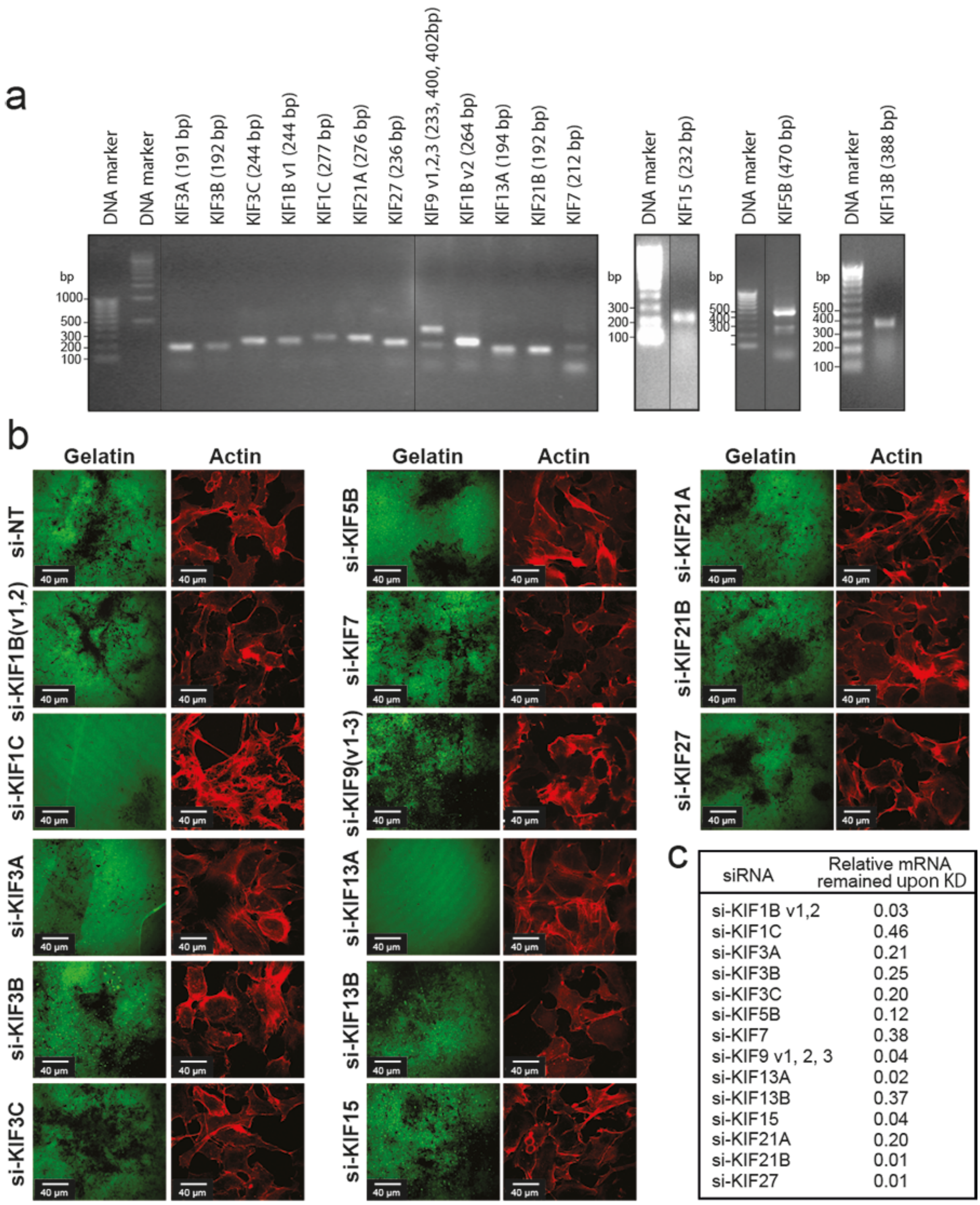


c

| siRNA | Relative mRNA remained upon KD |
|---|---|
| si-KIF1B v1,2 | 0.03 |
| si-KIF1C | 0.46 |
| si-KIF3A | 0.21 |
| si-KIF3B | 0.25 |
| si-KIF3C | 0.20 |
| si-KIF5B | 0.12 |
| si-KIF7 | 0.38 |
| si-KIF9 v1, 2, 3 | 0.04 |
| si-KIF13A | 0.02 |
| si-KIF13B | 0.37 |
| si-KIF15 | 0.04 |
| si-KIF21A | 0.20 |
| si-KIF21B | 0.01 |
| si-KIF27 | 0.01 |

**Figure S2. Screening KIFs involved in MT1-MMP vesicle trafficking for degradation of ECM substratum.**

**a**. Expression of KIF genes screened in HT-1080. The mRNA extracted from HT1080 was subjected to RT-PCR (30 cycles) for expression of KIF genes selected for screening.

**b**. HT-1080 cells were transfected with siRNA for selected KIFs and subjected to Alexa488-gelatin film degradation assay.

**c**. Relative mRNA remained upon knockdown (KD) compared to non-target siRNA (si-NT)-transfected cells.

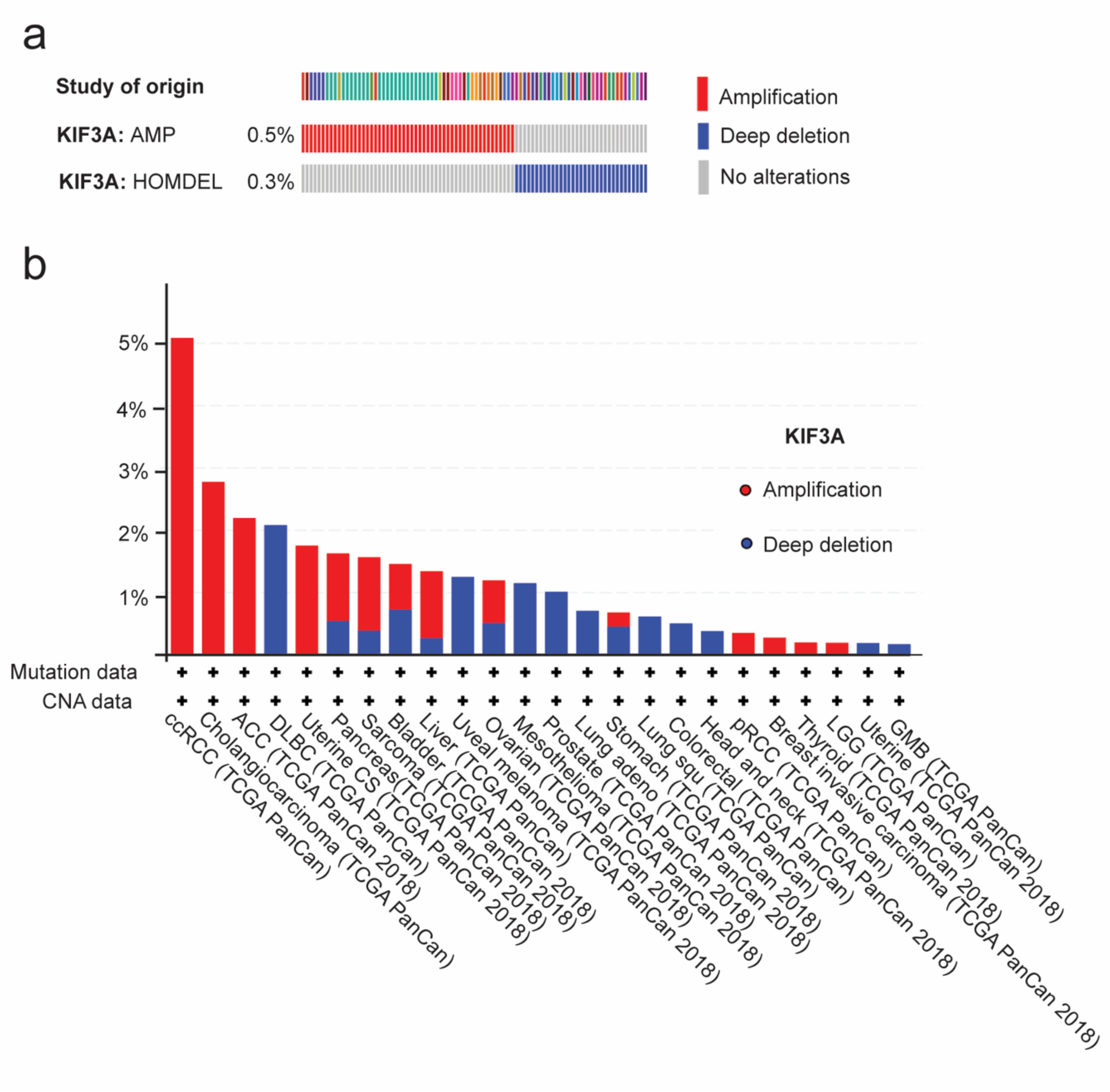


**Supplementary Figure S3. Genomic analysis of KIF3A.**

**a.** The oncoprint of KIF3A was identified. Genetic alterations of KIF3A. The columns represent patients from studies within TCGA and the rows gene alterations like amplification and deep deletions.

**b.** Genetic alterations of KIF3A summarised according to the study type. Amplifications are shown in red and deep deletions are blue.
